# The archaeal family 3 polyphosphate kinase reveals a function of polyphosphate as energy buffer under low energy charge

**DOI:** 10.1101/2024.08.28.610084

**Authors:** Svenja Höfmann, Christian Schmerling, Christina Stracke, Felix Niemeyer, Torsten Schaller, Jacky L. Snoep, Christopher Bräsen, Bettina Siebers

## Abstract

Inorganic polyphosphate, a linear polymer of orthophosphate residues linked by phosphoanhydride bonds, occurs in all three domains of life and plays a diverse and prominent role in metabolism and cellular regulation. While the polyphosphate metabolism and its physiological significance have been well studied in bacteria and eukaryotes including human, there are only few studies in archaea available so far. In Crenarchaeota including members of *Sulfolobaceae*, the presence of polyphosphate and degradation via exopolyphosphatase has been reported and there is some evidence for a functional role in metal ion chelation, biofilm formation, adhesion and motility, however, the nature of the crenarchaeal polyphosphate kinase is still unknown. Here we used the crenarchaeal model organism *Sulfolobus acidocaldarius* to study the enzymes involved in polyphosphate synthesis. The two genes annotated as thymidylate kinase (*saci_2019* and *saci_2020*), localized downstream of the exopolyphosphatase, were identified as the missing polyphosphate kinase in *S. acidocaldarius* (*Sa*PPK3). Thymidylate kinase activity was confirmed for Saci_0893. Notably Saci_2020 showed no polyphosphate kinase activity on its own but served as regulatory subunit (rPPK3) and was able to enhance polyphosphate kinase activity of the catalytically active subunit Saci_2019 (cPPK3). Heteromeric polyphosphate kinase activity is reversible and shows a clear preference for polyP-dependent nucleotide kinase activity, i.e. polyP-dependent formation of ATP from ADP (12.4 U/mg) and to a lower extent of GDP to GTP whereas AMP does not serve as substrate. PPK activity in the direction of ATP-dependent polyP synthesis is rather low (0.25 U/mg); GTP was not used as phosphoryl donor. A combined experimental modelling approach using quantitative ^31^P NMR allowed to follow the reversible enzyme reaction for both ATP and polyP synthesis. PolyP synthesis was only observed when the ATP/ADP ratio was kept high, using an ATP recycling system. In absence of such a recycling system, all incubations with polyP and PPK would reach an equilibrium state with an ATP/ADP ratio between 3 and 4, independent of the initial conditions. Structural and sequence comparisons as well as phylogenetic analysis reveal that the *S. acidocaldarius* PPK is a member of a new PPK family, named PPK3, within the thymidylate kinase family of the P-loop kinase superfamily, clearly separated from PPK2. Our studies show that polyP, in addition to its function as phosphate storage, has a special importance for the energy homeostasis of *S. acidocaldarius* and due to its reversibility serves as energy buffer under low energy charge enabling a quick response to changes in cellular demand.

## Introduction

Inorganic polyphosphate (polyP) is a linear polymer of three up to hundreds of orthophosphates linked by high-energy phosphoanhydride bonds (Kornberg, Rao et al. 1999). Among many function (described in detail below) polyP serves as an energy source for nucleotide (ATP) formation, as a reservoir of P_i_ essential for metabolism and cell growth (P_i_ homeostasis) and as an ATP substitute. Furthermore, its significance as a potential primordial energy currency in a pre-ATP world is under discussion (Kornberg 1995). PolyP and the enzymes involved in polyP metabolism will be key for future biotechnical applications of thermoacidophilies (Bacteria and Archaea) as engineered bio-factories, for industrial biomining (bioleaching) (Orell, Navarro et al. 2010), for enhanced biological phosphorus removal (EBPR, a treatment process by which microbes can remove excessive Pi from wastewater or sewage sludge (Blank 2023, Demling, Baier et al. 2024)) and for the use in biocatalysis and metabolic engineering, such as serving as an energy (ATP) regeneration system or facilitating the production of (modified) nucleoside triphosphates (NTPs) (Andexer and Richter 2015, Keppler, Moser et al. 2022, Yang, Zhu et al. 2023). In addition, polyP is used as food additive in the meat and dairy industry to improve flavor, water binding, color retention and emulsification, while retarding oxidative rancidity and antimicrobial activity against Bacteria, with gram-positive Bacteria being more sensitive, has been reported (for review see (Rao, Gómez-García et al. 2009)).

PolyP was first isolated from yeast (Liebermann 1888) and then identified as dense, metachromatic granules (so-called volutins) in many microorganisms (Kornberg 1995). Today, polyP is known to occur in all three domains of life, including Archaea (Rao, Gómez-García et al. 2009). Despite the ubiquitous distribution of polyP, research on the metabolism and functions of “the forgotten polymer” (Kornberg 1995) was scarce until the discovery of polyP kinase (PPK), the enzyme involved in polyP synthesis, in *Escherichia coli* (Ahn and Kornberg 1990). Since then, the metabolism of polyP has been studied and started to elucidate the main routes and enzymes involved in polyP synthesis and degradation in many bacteria and eukaryotes and has revealed diverse physiological roles (for review see (Seufferheld, Alvarez et al. 2008, Rao, Gómez-García et al. 2009, Neville, Roberge et al. 2022). However, the metabolism of polyP in Archaea remains mainly unexplored.

Polyphosphate kinases (PPKs) are reversible and catalyze polyP synthesis from nucleotides (NTPs) in the presence of divalent metal ions and polyP degradation for nucleotide formation in the presence of NDPs or NMPs. Two different families of PPKs (EC 2.7.4.1) involved in polyP metabolism have been described: PPK1 catalyze the reversible transfers of the terminal γ-phosphate from ATP to polyP, with clear preference for polyP synthesis (Ahn and Kornberg 1990). PPK1 homologs are found in many Bacteria and few eukaryotes such as the social slime mold *Dictyostelium discoideum* (Zhang, Gómez-García et al. 2007). PPK2s are also capable of synthesizing polyP, but preferentially catalyze the reverse reaction, the phosphorylation of nucleotide mono- or diphosphates, and thus function as polyP-dependent nucleotide kinases (Motomura, Hirota et al. 2014) (for review see (Neville, Roberge et al. 2022)). PPK2s do not share any sequence similarity to PPK1 and phylogenetic, structural and sequence analysis suggests that they evolved from P-loop kinases and possess a common evolutionary origin with bacterial thymidylate kinases (Nocek, Kochinyan et al. 2008). PPK2s exhibit diverse substrate specificity and metal cofactor preferences and were categorized into three classes based on their nucleoside phosphate preference (Ishige, Zhang et al. 2002, Motomura, Hirota et al. 2014)): The class I PPK2s with only one PPK domain utilize polyP to phosphorylate nucleoside diphosphates mainly ADP or GDP (Nocek, Kochinyan et al. 2008). The class II PPK2s with either one or two PPK domains catalyze the phosphorylation of nucleoside monophosphates e.g. AMP to ADP by polyP (poly-AMP phosphotransferase) (Nocek, Kochinyan et al. 2008). Finally, the class III PPK2s with one PPK domain either phosphorylate nucleotide mono- or diphosphates and thus are able to synthesize the direct formation of ATP from AMP and polyP (Motomura, Hirota et al. 2014). The distribution of PPK1s and PPK2s in the bacteria is very diverse. While some organisms have only one PPK1 or PPK2, some possess multiple homologues and others have both PPK1 and PPK2 (Rao, Gómez-García et al. 2009, Neville, Roberge et al. 2022). For example, whereas *Escherichia coli* possesses only one ppk1 gene, *Ralstonia eutropha* comprises two *ppk1* genes and five *ppk2* genes (Rosigkeit, Kneißle et al. 2021).

Despite the widespread presence of PPK1 and PPK2, the mechanisms for polyP synthesis are not universally conserved and in many eukaryotes the synthesis mechanism remains unknown. In yeast, polyP is synthesized by a vacuolar transporter chaperone 4 (VTC4) complex, which has also been identified in other eukaryotes such as *Trypanosoma brucei* (Guan, Chen et al. 2023). The amoeba/social slime mold *Dictyostelium discoideum* employs, in addition to a bacterial PPK1, an actin-related protein (Arps) complex that synthesizes an actin-like filament in conjunction with polyP in a fully reversible reaction (Gómez-García and Kornberg 2004, Zhang, Gómez-García et al. 2007).

The most common enzyme for the degradation of polyP besides PPK found in all three domains of life is expopolyphosphatase (PPX), which hydrolyzes the terminal phosphoanhydride bond in the linear polymer and releases P_i_ at the expense of energy. In addition, endopolyphosphatases (PPNs) that cleave inside the polymer forming polyPs of different chain length has been reported from yeast or animal cells (Sethuraman, Rao et al. 2001, Rao, Gómez-García et al. 2009, Blank 2012). Last but not least, several enzymes that employ both, either polyP or ATP, as phosphate donor such as NAD kinase and glucokinase have been described (Rao, Gómez-García et al. 2009) also from archaea (Sakuraba, Kawakami et al. 2005).

PolyP acts as a versatile regulator influencing cellular processes, including for example stress response, bacterial homeostasis, motility, biofilm formation, quorum sensing, genetic competence, virulence, DNA replication, and cytokinesis (for review see (Seufferheld, Alvarez et al. 2008, Rao, Gómez-García et al. 2009, Xie and Jakob 2019, Neville, Roberge et al. 2022). In addition, polyP serves as channel for DNA entering (genetic competence), and as inorganic polyanion functions as a metal ion chelator, buffer against alkali ions, as well as primordial chaperone and pro-aggregation molecule (Xie and Jakob 2019). In eukaryotes, polyP exhibits varied cellular localizations and impacts processes like DNA synthesis and Pi homeostasis in yeast. Mammalian studies associate polyP chain length and concentration with bone healing, nerve transduction, blood clotting, and cancer metastasis (Xie and Jakob 2019). Finally, a non-covalent post translational modification (PTM) by polyphosphorylation of internal lysine and consecutive histidine residues was observed in yeast, which regulates protein-protein interactions and enzyme activity (Azevedo, Livermore et al. 2015, Neville, Lehotsky et al. 2023, Neville, Lehotsky et al. 2024)

Unlike bacteria and eukaryotes, our knowledge of polyphosphate (polyP) metabolism in archaea is limited. Although polyP’s presence in archaea was recognized in 1983, its functions remain relatively unexplored (Scherer and Bochem 1983). The presence of polyP in these microbes was already described in 1983 (Scherer and Bochem 1983) and polyP granules are known to be present in several archaea, including *Sulfuracidifex* (formerly *Sulfolobus*) *metallicus*, *Metallospaera sedula*, *Saccharolobus* (formerly *Sulfolobus*) *solfataricus* and *S. acidocaldarius* (Remonsellez, Orell et al. 2006, Orell, Navarro et al. 2012). Furthermore, some studies have started to elucidate some of the functional roles mediated by polyP in Archaea. In the Euryarchaeon/methanogen *Methanosarcina mazei* P_i_ starvation/surplus experiments revealed induction of *ppk1* gene transcripts and subsequent polyP accumulation (Paula, Chin et al. 2019). In *S. metallicus* and *S. solfataricus*, the influence of polyP accumulation in response to metal toxicity (e.g. copper) was studied due to its role in bioleaching (Remonsellez, Orell et al. 2006, Rivero, Torres-Paris et al. 2018, Soto, Recalde et al. 2019). Furthermore, in *S. acidocaldarius*, a reduction in polyP by *Sa*PPX overexpression has been linked to decreased motility, adhesion and biofilm formation (Recalde, van Wolferen et al. 2021). However, some of the most important enzymes involved in polyP metabolism in many archaea – such as PPKs and PPXs – have so far not been identified and/or have not been biochemically characterized.

Notably, while bioinformatic analyses in extremophiles revealed the presence of key enzymes for polyP synthesis and degradation in all extremophilic bacteria, many extremophilic archaea encode incomplete sets of enzymes involved in polyP metabolism. For example, although many methanogenic archaea seem to encode a complete polyP pathway, many halophilic Euryarchaeota (i.e. Haloferacales, Natrialbales and Halobacteriales) lack genes encoding catabolic PPX, whereas all Crenarchaeota lack genes encoding homologs for reversible polyP synthesis via PPKs (e.g. PPK1s, PPK2, Arps) (Orell, Navarro et al. 2012, Paula, Chin et al. 2019, Wang, Liu et al. 2019). Therefore, although there is good evidence for polyP formation in *Sulfolobales* (and Crenarchaeota more generally) (Orell, Navarro et al. 2012), the enzymes involved in polyP synthesis in these organisms remain unknown. Importantly, overexpression of PPX in *Sulfolobales* led to reduced polyP levels and compromised cellular functions, suggesting that this is one of the enzymes involved in polyP degradation in these archaea (Recalde, van Wolferen et al. 2021). Here we report the identification and characterization of the missing PPK in the crenarchaeal model *S. acidocaldarius*, thriving at 76 °C and pH 2-3 (Brock, Brock et al. 1972).

## Results

### Polyphosphate synthesis in *S. acidocaldarius*

Although there is good evidence for polyP formation and PPX function in Sulfolobales (and Crenarchaeota more generally) the enzymes involved in polyP synthesis in these organisms remain unknown. Despite a lack of bacterial/eukaryotic PPK homologues (Orell, Navarro et al. 2012, Paula, Chin et al. 2019, Wang, Liu et al. 2019), closer inspection of the gene neighborhood of the *ppx* gene (*saci_2018*) revealed a conserved gene organization resembling the one in Bacteria and other Archaea, although classical PPK1 and/or PPK2 enzymes were missing. Available transcriptome data confirmed that the *ppx* gene forms an operon with genes encoding a phosphohistidine phosphatase/mutase SixA (*saci_2017*), two thymidylate kinases (dTMPKs, *saci_2019* and *saci_2020*) as well as a ’conserved histidine α-helical domains, CHAD domains’ CHAD-domain containing protein (*saci_2021*) (Fig. 1A, SI Fig. 1). A third dTMPK-encoding gene was identified at a different locus in the *S. acidocaldarius genome* (*saci_0839*). Notably, dTMPKs have been characterized as the closest structural homologs of PPK2 (Leipe, Koonin et al. 2003, Nocek, Kochinyan et al. 2008, Motomura, Hirota et al. 2014), suggesting that these enzymes may serve as the PPK enzyme in *S. acidocaldarius.* Consequently, all three proteins were recombinantly expressed and their respective enzymatic activities were examined.

**Figure 1.**
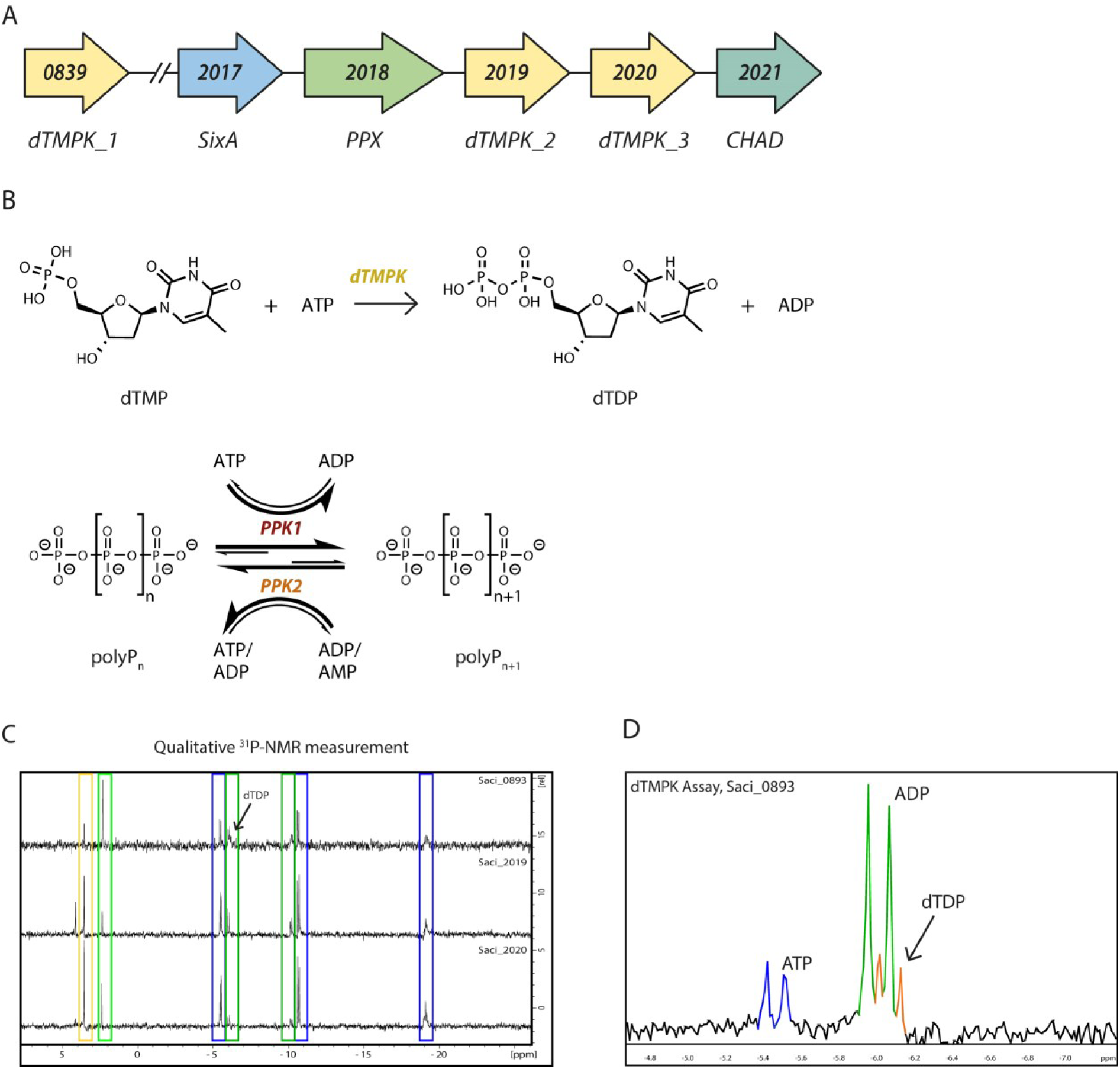
Investigation of polyphosphate kinase and thymidylate kinase candidates in *S. acidocaldarius*. (A) Localization of annotated dTMPKs in the *S. acidocaldarius* genome. Two thymidylate kinase genes *saci_2019* (dTMPK_2) and *saci_2020* (dTMPK_3) are located adjacent to the *ppx* gene (*saci_2018*) while the third candidate is separate in the genome (*saci_0893*, dTMPK_1). In addition, two genes encoding SixA and CHAD domain protein homologues are located in the conserved polyP operon (see also SI Fig. 1). (B) Enzyme reactions catalyzed by dTMPKs and PPKs. (C) ^31^P NMR spectra of dTMPK assays of the recombinant Saci_0893, Saci_2019 and Saci_2020, respectively. The assays were performed at 70 °C for 8 hours in presence of 2 mM ATP, 1 mM dTMP, 4 mM MgCl_2_, 50 mM KCl with 10 µg/ml of the respective enzyme in 50 mM TRIS/HCl pH 7.4 with 10% (v/v) D_2_O (total volume 1 ml). The respective signals for ATP (blue), ADP (green), dTMP (yellow) and P_i_ (light green) are marked. (D) Magnification showing the formation of dTDP in the presence of Saci_0893. The dTDP signal (orange) overlaps with one ADP signal (green).

### Cloning, expression, purification and acitvity of the three annotated thymidylate kinases (Saci_0893, Saci_2019, Saci_2020)

The genes encoding the three annotated thymidylate kinases were cloned into the expression vectors pET15b (*saci_0893*, *saci_2019*) and pET28b (*saci_2020*) for heterologous expression in *E. coli* Rosetta. The N-terminal His-tagged proteins were purified using IMAC and a total of 200 µg of Saci_0893, 250 µg of Saci_2019, and 120 µg of Saci_2020 were obtained from 1.0 g, 6.0 g, and 9.0 g of *E. coli* cells (wet weight), respectively (SI Fig. 2). Thymidylate kinases (EC: 2.7.4.9, dTMPK) catalyze the Mg^2+^-dependent phosphorylation of deoxythymidine monophosphate (dTMP) to deoxythymidine diphosphate (dTDP) using ATP as a phosphoryl donor (Fig. 1B). The dTMPK activities of the three enzymes were qualitatively analyzed using ^31^P NMR spectroscopy (Fig. 1C). Thymidylate kinase activity was tested with dTMP (1 mM) as the substrate and ATP (2 mM) as the phosphoryl donor. Whereas Saci_2020 showed no activity, Saci_0893 and Saci_2019 showed ADP formation from ATP, however, only Saci_0893 showed ATP-dependent formation of dTDP from dTMP (Fig. 1D).

To analyze for a potential PPK function of the three dTMPKs we performed initial experiments (discontinuous assays) by monitoring the ATP-dependent formation of polyP, ADP, and P_i_ over time using the MicroMolar Polyphosphate Assay Kit (ProFoldin), the PK-LDH assay, and the malachite green assay, respectively (SI Fig. 3). Our results showed that only Saci_0893 possessed dTMPK activity with additional formation of ADP in presence of the substrate dTMP (1mM), whereas Saci_2019 displayed PPK activity forming polyP starting from ATP, without requiring a precursor. In contrast, Saci_2020 showed no evidence of dTMPK or PPK activity. The specific activity of dTMPK Saci_0893 was determined using a continuous coupled PK LDH assay at 55 °C following ADP formation, which revealed a dTMP-dependent specific activity of of 4.6 U/mg (V_max_) and K_M(dTMP)_ of 0.013 mM (SI Fig. 4).

Since a two-domain structure with an inactive N-terminal domain has been reported for class II PPK2 from *P. aeruginosa* (Nocek, Kochinyan et al. 2008), we examined the influence of Saci_2020 on the PPK activity of Saci_2019. In a series of experiments, we fixed the amount of Saci_2019 to 10 µg/ml and added varying amounts of Saci_2020 and determined the specific PPK activity based on ADP formation at 70°C (discontinuous PK-LDH assay) and observed a steady increase in PPK activity with increasing amounts of Saci_2020 (SI Fig.5A). To determine the optimal molar ratio of the two enzymes a titration experiment was performed at 70°C (molar ratios from 10:0 to 0:10 (Saci_2019 to Saci_2020)) by following polyP and ADP formation (discontinuous polyP and PK-LDH assay) (SI Fig. 5B). The highest PPK activity was observed at a molar ratio of 7:3–5:5, with a specific activity of 329 mU/mg protein. (PK-LDH assay, 139 mU/mg polyP assay) Therefore, for all further PPK experiments with the heteromeric enzyme, we used a 1:1 molar ratio of Saci_2019 to Saci_2020. Hereafter the enzyme is named *Sa*PPK3 (heteromeric enzyme), whereas the enzyme subunits are called cPPK3 for the catalytically active subunit (Saci_2019) and rPPK3 for the regulatory subunit (Saci_2020). The heteromeric *Sa*PPK3 (Saci_2019 and Saci_2020,1:1 ratio) showed highest activity at neutral pH (pH 7) and 70°C with only 12% residual activity at 80°C (SI Fig. 6). *Sa*PPK3 activity was dependent on Mg^2+^ and no activity was observed in absence of metal ions (data not shown). Gelfiltration experiments revealed a monomeric structure for cPPK3 (Saci_2019) with higher aggregation states and a mixture of monomers and tetramers for the heteromeric *Sa*PPK3 (Saci_2019 and Saci_2020 (1:1)) suggesting that the complex might either dissolve on the column or the *in vitro* complex formation is not complete (SI Fig. 7).

For an initial characterization of the heteromeric *Sa*PPK3 the discontinuous polyP assay was used and a specific activity (Vmax) of 63.3 mU/mg and K_M(ATP)_ of 0.95 mM were determined (SI Fig. 8). Notably, a strong inhibition of the reactions at ATP concentration above 2 mM was observed and to determine if ADP formation might influence the enzyme activity of heteromeric *Sa*PPK3, polyP formation (1.5 mM ATP) was monitored over time in the presence and absence of ADP (2 mM). The addition of ADP resulted in the complete inhibition of polyP synthesis (SI Fig. 9), and therefore we next examined the reversibility of the PPK reaction.

In initial experiments we determined the reversibility of the PPK reaction by examining ATP or GTP formation from ADP and GDP in the presence of polyP (50 µM polyP_45_) as a phosphate donor. Initial enzyme assays were conducted using 5 µg/ml of heteromeric *Sa*PPK3 (SI Fig. 10A (ADP), B (GDP)) and the single subunits cPPK3 (Saci_2019) and rPPK3 (Saci_2020) (SI Fig. 10C, D (both with ADP)). PolyP was utilized, and ATP or GTP formation at 70°C was monitored using the discontinuous polyP and hexokinase (HK)-glucose-6-phosphate dehydrogenase (G6PDH) assays. The heteromeric *Sa*PPK3 showed a specific activity with ADP as P_i_ donor of 56.3 mU/mg (polyP detection) and 1.40 U/mg (ATP detection). For GDP the activity was significantly reduced (6.13 mU/mg (polyP detection) and 0.455 U/mg (GTP detection)). For the homomeric subunits using ADP as P_i_ acceptor, only cPPK3 (Saci_2019) catalyzed ATP synthesis, with a specific activity of 148 mU/mg, which was about 10-times lower than that of the heteromeric *Sa*PPK3 (SI Fig. 10C, D). Notably, due to the low activity no decrease in the polyP concentration was observed for cPPK3. rPPK3 (Saci_2020) showed no activity at all.

### Kinetic characterization of *Sa*PPK3, model construction and validation

To address the kinetic and regulatory properties of *Sa*PPK3 and PPK3 we used our established iterative experimental and modelling approach (Shen, Kohlhaas et al. 2020, Kuschmierz, Shen et al. 2022) and first determined the initial rate kinetics (Fig. 2).

**Figure 2.**
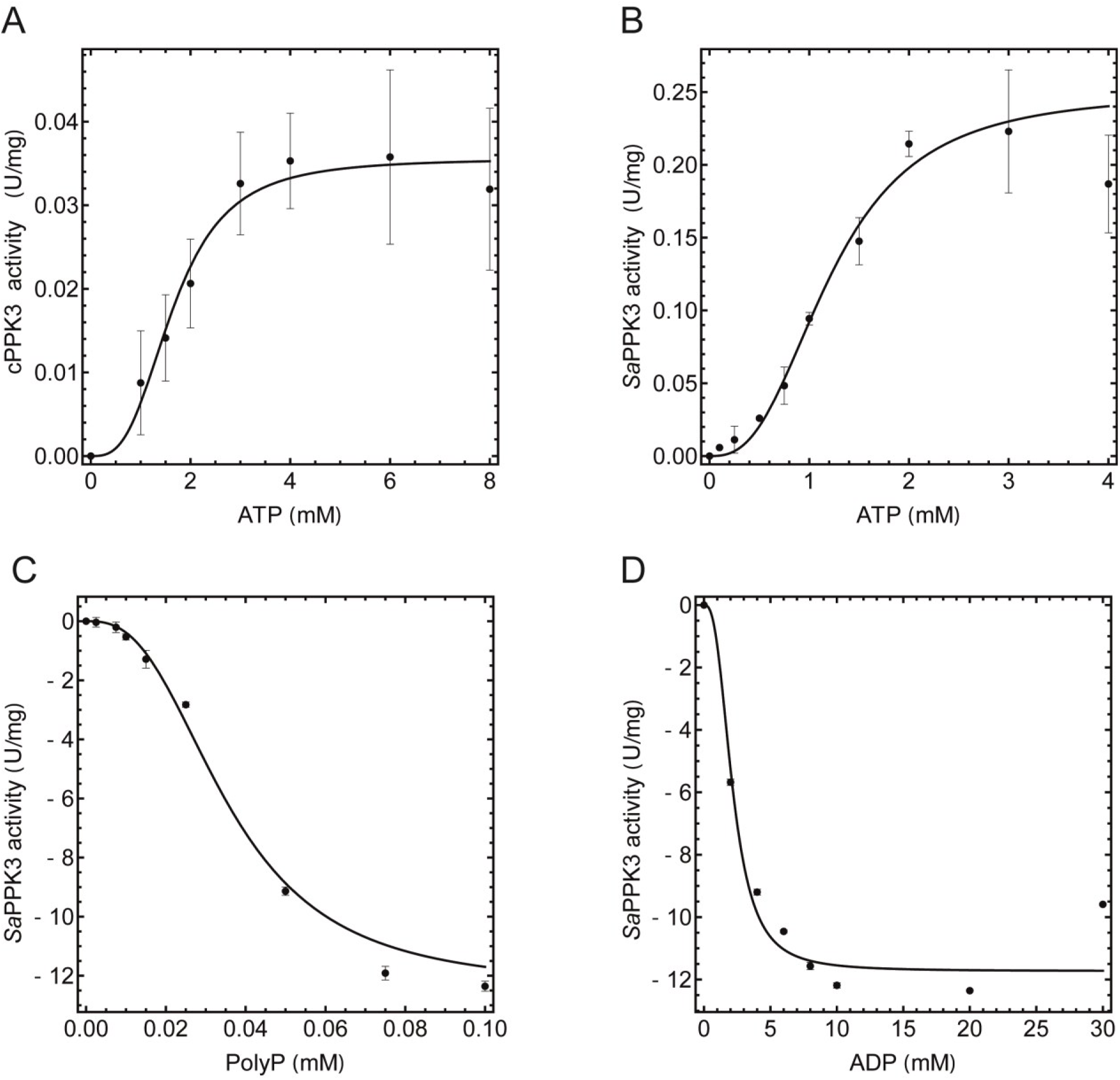
Initial rate kinetics of the recombinant, homomeric cPPK3 (Saci_2019) and heteromeric *Sa*PPK3 (Saci_2019 and Saci_2020). The kinetic properties of the recombinant, homomeric PPK3 (Saci_2019) (A) and the heteromeric *Sa*PPK3 (Saci_2019 and Saci_2020, (1:1) (B) with respect to ATP-dependent polyP synthesis were determined by following ADP formation at 70°C via the discontinuous PK-LDH assay. The polyP-dependent nucleotide kinase activity, i.e. ATP formation from polyP of the heteromeric *Sa*PPK3 with different polyP (C) or ADP concentrations (D) was determined at 70°C via the discontinuous HK-G6PDH assay. For all experiments three independent measurements (n=3) were performed and error bars indicate the standard error of the mean (SEM).

A kinetic model for polyP metabolism was constructed based on initial rate kinetics for the PKK, and degradation kinetics for polyP and PEP. The model had to be adapted in several ways to simulate the polyP dynamics in the incubation assays (see material and methods). The rate equations for the reactions are shown in eqs. 1-5, with the fitted parameters in table 1.

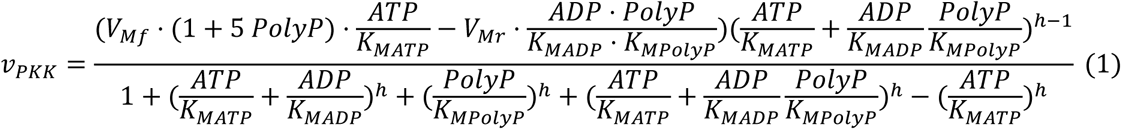

**Table 1:**
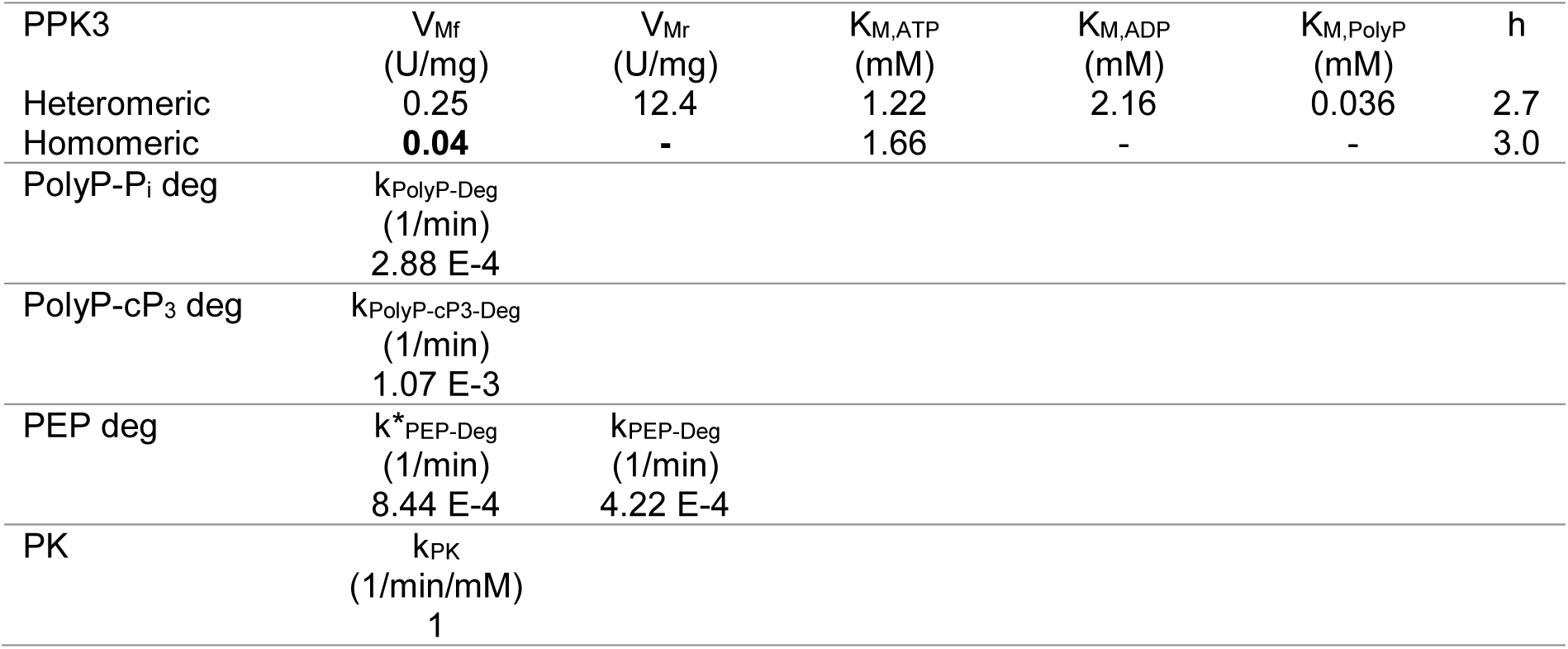
Model parameters for PPK, degradation and ATP recycling reactions

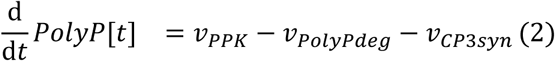

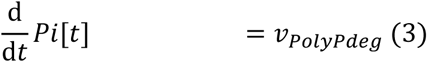

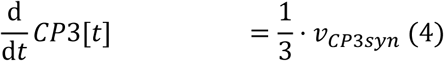

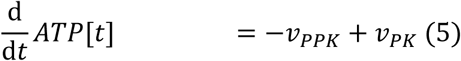

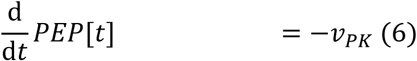

#### PolyP formation

The kinetic characterization of the homomeric cPPK3 (Saci_2019) with ATP as substrate (following ADP formation at 70°C via the discontinuous PK-LDH assay) showed sigmoidal saturation kinetics, resulting in a V_max_ of 0.04 U/mg and an apparent K_M(ATP)_ of 1.66 mM (70 °C) (Fig. 2A). For the heteromeric *Sa*PPK3 (Fig. 2B), a specific activity of 0.25 U/mg and a K_M(ATP)_ of 1.22 mM were estimated by fitting a Hill equation (h=2.71) to the data. Notably, some deviation in V_max_-values between the different enzyme isolations were observed. Despite this, the heteromeric *Sa*PPK3 displayed an approximately 6-fold higher specific activity than the homomeric cPPK3. No enzyme activity was detected when GTP was used as the substrate (data not shown).

#### Nucleotide formation

The kinetic properties of the heteromeric *Sa*PPK3 enzyme were studied using polyP_45_ and ADP as the (co-)substrates (Fig. 2C, D). The results showed that the enzyme had a higher catalytic efficiency for polyP-dependent nucleotide formation, with the following kinetic parameters obtained by fitting a Hill equation: V_max_-value of 12.4 U/mg, K_M_-value of 0.036 mM for polyP45 and 2.16 mM for ADP. Note that differences in specific activity between different isolations made it necessary to normalize the activities to the isolation with the higher activity such that a single equation could be fitted to the total data set (Fig. 2C, D). This indicated that the enzyme had a clear preference (50-fold higher specific activity) for polyP-dependent nucleotide formation.

To follow polyP synthesis and degradation, linked to ATP and ADP dynamics, we used quantitative ^31^P NMR analysis (Fig. 3, corresponding NMR spectra SI Fig. 11 and 12). First, we had to test the stability of polyP_45_ and of phosphoenolpyruvate (PEP, used for ATP recycling) at the high temperatures used during the incubation (Fig. 3 A-C). PolyP was degraded to an unknown phosphate compound identified as cyclic-triphosphate (cP3) (singlet at -20.5 ppm) and Pi (SI Fig. 13) with rate constants of 1*10^-3^ 1/min and 0.3*10^-3^ 1/min respectively, while the degradation of PEP was found to be dependent on the presence of PPK (0.4*10^-3^ 1/min in absence and 0.8*10^-3^ 1/min in presence of PPK) (Fig. 3 A, B). The non-enzymatic formation of cP_3_ from polyP at high temperature in the presence of metal ions has been reported previously (Rulliere, Perenes et al. 2012). Further *in vitro* studies using ^31^P NMR spectroscopy confirmed the increase of non-enzymatic cP_3_ and P_i_ formation from polyP in presence of divalent metal ions (MgSO_4_, 2.5 mM, 6-fold) and monovalent ions (KH_2_PO_4_, 500 µM, 2-fold) at 70 °C (SI Fig. 13). cP_3_ was stable under the chosen assay conditions and time-dependent hydrolysis to orthophosphate was not observed.

**Figure 3.**
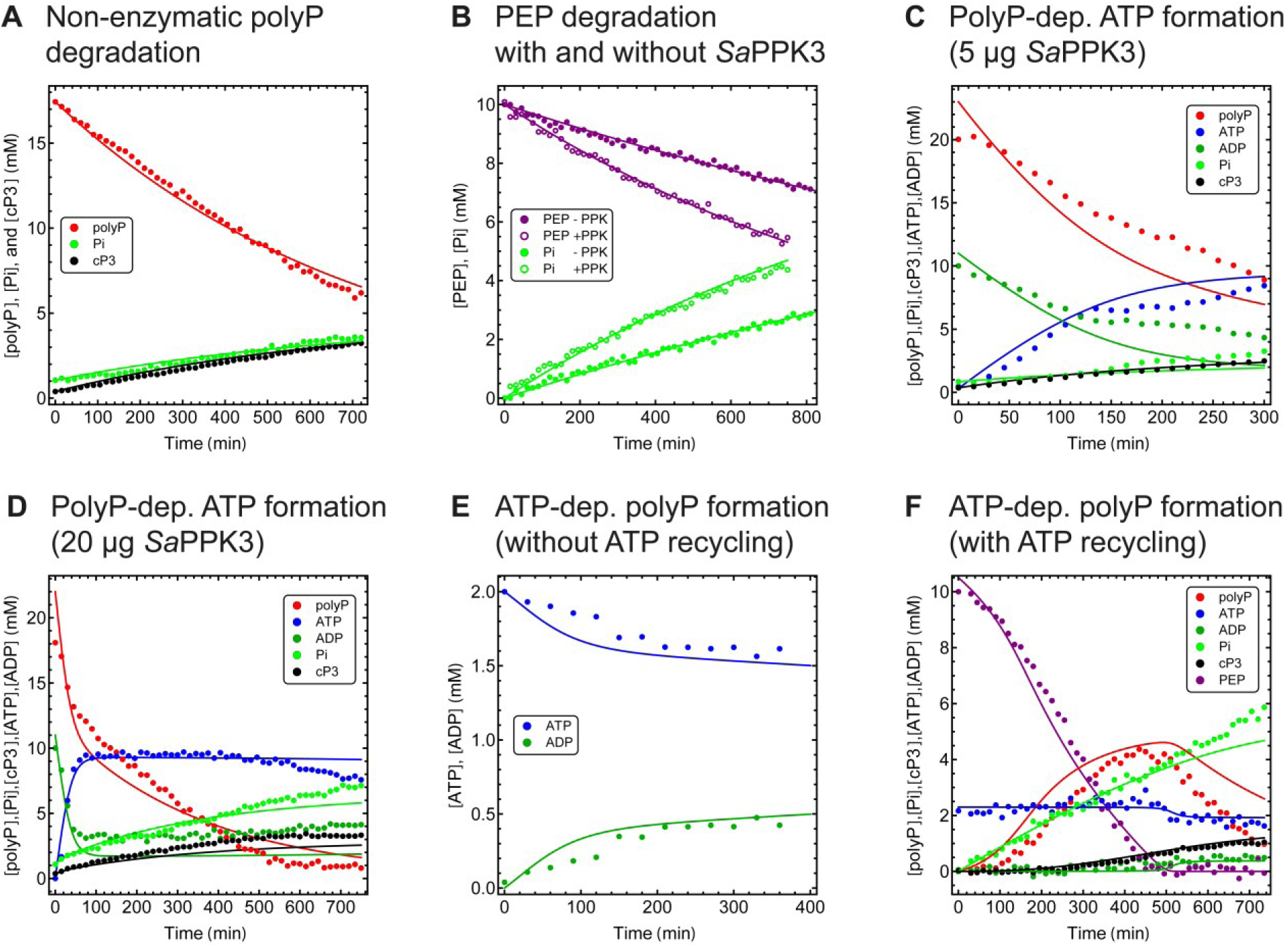
Analysis of the reversible heteromeric *Sa*PPK3 using ^31^P NMR spectroscopy. The non-enzymatic degradation of polyP to cP_3_ and P_i_ (polyP_45_ hydrolysis) (A), PEP hydrolysis in absence and presence of *Sa*PPK3 (B), the polyP-dependent ATP formation with 5 µg *Sa*PPK3 (C) and 20 µg/ml (D) and the ATP-dependent polyP formation without (E) and with ATP recycling system (F) (both 20 µg/ml *Sa*PPK3) is shown. The concentration of polyP is expressed in mM P_i_. The non-enzymatic polyP hydrolysis (A) was studied using the assay conditions for polyP-dependent ATP formation (see C, D) (10 mM ADP, 10 mM MgCl_2_, 0.5 mM polyP_45_) but without heteromeric *Sa*PPK3. PEP hydrolysis (B) was followed using the assay conditions for ATP-dependent polyP formation (4 mM MgCl_2_, without ATP and *Sso*PK) in presence and absence of 20 µg/ml heteromeric *Sa*PPK3. ATP formation from poylP (C, D) was assayed with 10 mM ADP, 10 mM MgCl_2_, 0.5 mM polyP_45_ and 5 µg/ml (C) or 20 µg/ml (D) of heteromeric *Sa*PPK3. PolyP synthesis from ATP (E and F) was carried out with 2 mM ATP, 4 mM MgCl_2_ and 20 µg/ml heteromeric *Sa*PPK3 in absence (E) or presence of an ATP recycling system (8 µg/ml *Sso*PK, 10 mM PEP and 20 mM MgCl_2_ (final concentration) (F). All assays were performed at 70°C. The obtained signals in the ^31^P NMR spectra for the different phospho-compounds are shown (data points, for color code see figure, (see also SI Fig. 11, 12)). A mathematical model based on the non-enzymatic degradation of polyP to P_i_ and cP3, and the determined initial rate kinetics of *Sa*PPK3 were used to describe the experiment and model simulations which are shown in solid lines with corresponding colours to the experimental symbols.

To test the capacity of polyP and *Sa*PPK3 to act as a buffer for maintaining a constant ATP/ADP ratio (polyP-dependent nucleotide kinase activity), we started with an incubation of polyP (0.5 mM), *Sa*PPK3 (20 ug/ml) and ADP (10 mM). PolyP was rapidly dephosphorylated coupled to ATP synthesis until a stable ATP/ADP ratio of 3-4 was reached after 75 min (Fig. 3D, SI Fig.12A). Subsequently polyP was dephosphorylated to P_i_ and cP_3_ uncoupled from ATP formation.

The kinetic model, based on the initial rate kinetics, with small adaptations (see above) for *Sa*PPK3 and temperature stability of the phosphorylated intermediates could describe the different polyP incubations precisely. The rapid initial polyP dephosphorylation (−0.31 mM P_i_/min) is mostly driven by the *Sa*PPK3 (80%) but slows down strongly to about 10% of the initial rate with now all phosphate ending up in cP_3_ (58%) and P_i_ (42%) (Fig. 3D). A second incubation with the same initial conditions, but lower *Sa*PPK3 concentration was equally well described by the model, when set to the PKK concentration used in the incubation (5 ug/ml) (Fig. 2C, SI Fig. 12B).

We subsequently investigated the synthesis of polyP using ATP (Fig. 3E, SI Fig. 12C). To initiate the reaction, we incubated 2 mM ATP and 4 mM Mg^2+^ with 10% (v/v) D_2_O and 20 µg/ml of heteromeric *Sa*PPK3 at 70°C. ATP was converted to ADP until the equilibrium ATP/ADP ratio was reached (3-4) similar as observed in Fig. 3C, D) No polyP was detected in the incubation, while the model simulation for the experiment predicted a low polyP concentration of maximally 0.45 mM P_i_ (probably under the detection level for the NMR) at 200 min which was subsequently degraded. Thus, the rapid buildup of ADP strongly inhibited the *Sa*PPK3 thereby preventing significant production of polyP. To test this hypothesis, we subsequently included an enzymatic ATP recycling system based on recombinant pyruvate kinase from *S. solfataricus* (*Sso*PK; 20 mM MgCl_2_, 10 mM PEP, and 8 µg/ml *Sso*PK) (Fig. 3F, SI Fig. 12D).

During the incubation the PK-recycling system kept the ATP concentration close to 1 mM, with very low ADP concentrations, and the *Sa*PPK3 (20 µg/ml) showed significant and increasing activity over time. The kinetic model correctly described the dynamic behavior, albeit with a slight overprediction of the polyP concentration. As soon as the PEP runs out at about 500 min, the ATP/ADP ratio comes down to the equilibrium ratio, polyP synthesis stops and polyP is degraded to P_i_ and cP3. The relative efficiency and contribution of reactions to synthesis and degradation of polyP were similar to what was observed in the other incubations.

### Structural and phylogenetic analyses of PPK3 enzymes

Structural and sequence comparisons revealed that the *S. acidocaldarius* Saci_2019 cPPK3 and Saci_2020 rPPK3 adopt a dTMPK fold (Fig. 4) rather than a fold similar to PPK2s (Fig. 5). PPK2 enzymes contain an N-terminal extension forming an extended α helix and also differ with respect to the distal helix bundle behind the central five stranded β sheet shown in the right part of the structures depicted in Fig. 5.

**Figure 4.**
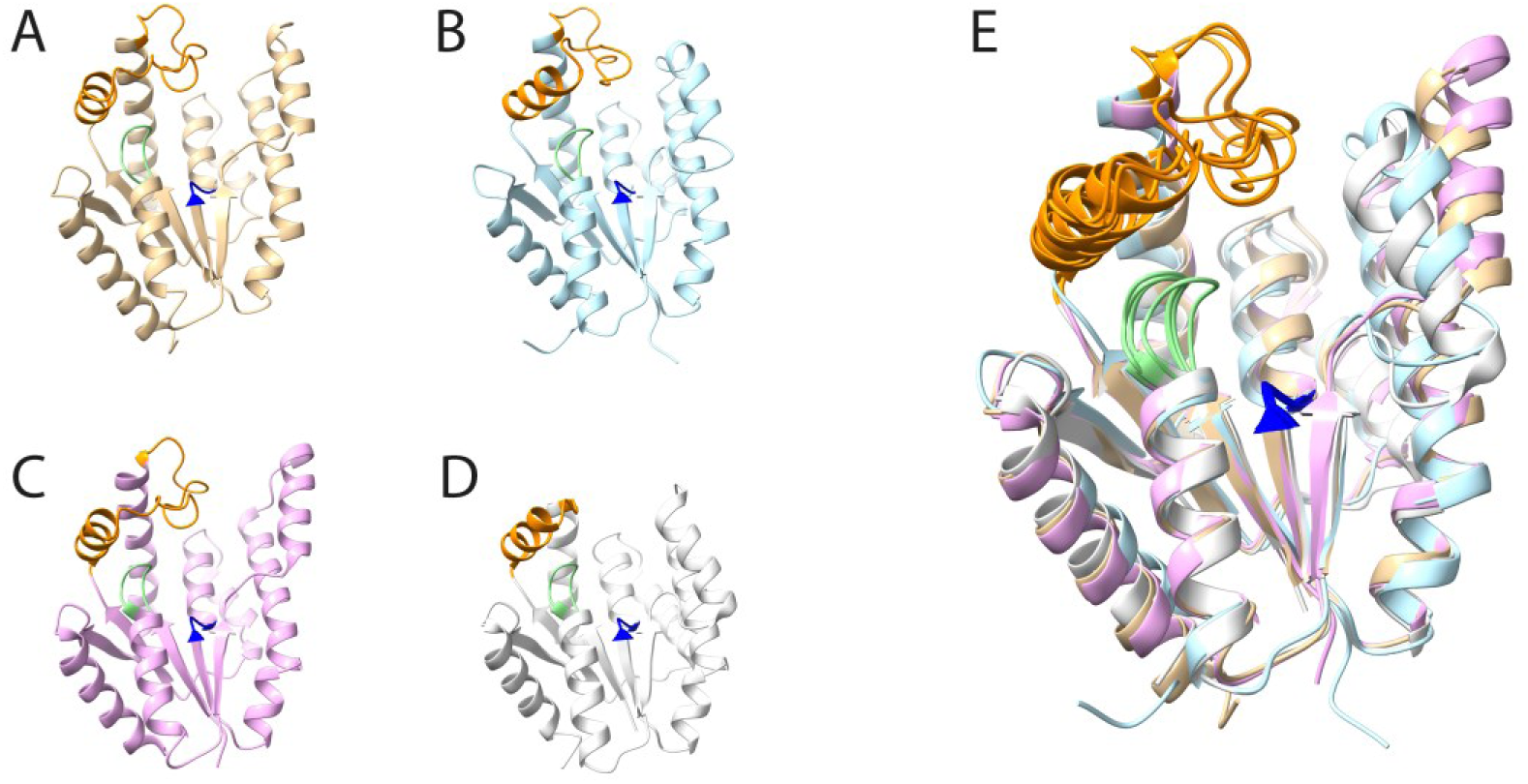
Structural comparison of the *S. acidocaldarius* cPPK3 (Saci_2019) and rPPK3 (Saci_2020) with other thymidylate kinase like proteins. The ribbon representations of the monomers of Saci_2019 (brown) (A) and Saci_2020 (blue) (B) (both models from the alphafold database) are comparatively illustrated with the crystal structure of the *Sulfurisphaera tokodaii* cPPK3 homologue STK_15430 (purple, pdb 5h70) (Biswas, Shukla et al. 2017) (C) and the thymidylate kinase from *Aquifex aeolicus* (grey, pdb 5xb2) (Biswas, Shukla et al. 2017) (D). The superimposition of all four monomers shown in (E) clearly shows that both, cPPK3 and rPPK3 from *S. acidocaldarius* adopt the typical thymidylate kinase fold (the structural motifs characteristic to the P-loop NTPases, i.e. the Lid region, the P-loop and the DRX motif (Leipe, Koonin et al. 2003), are highlighted in orange, green and blue, respectively).

**Figure 5.**
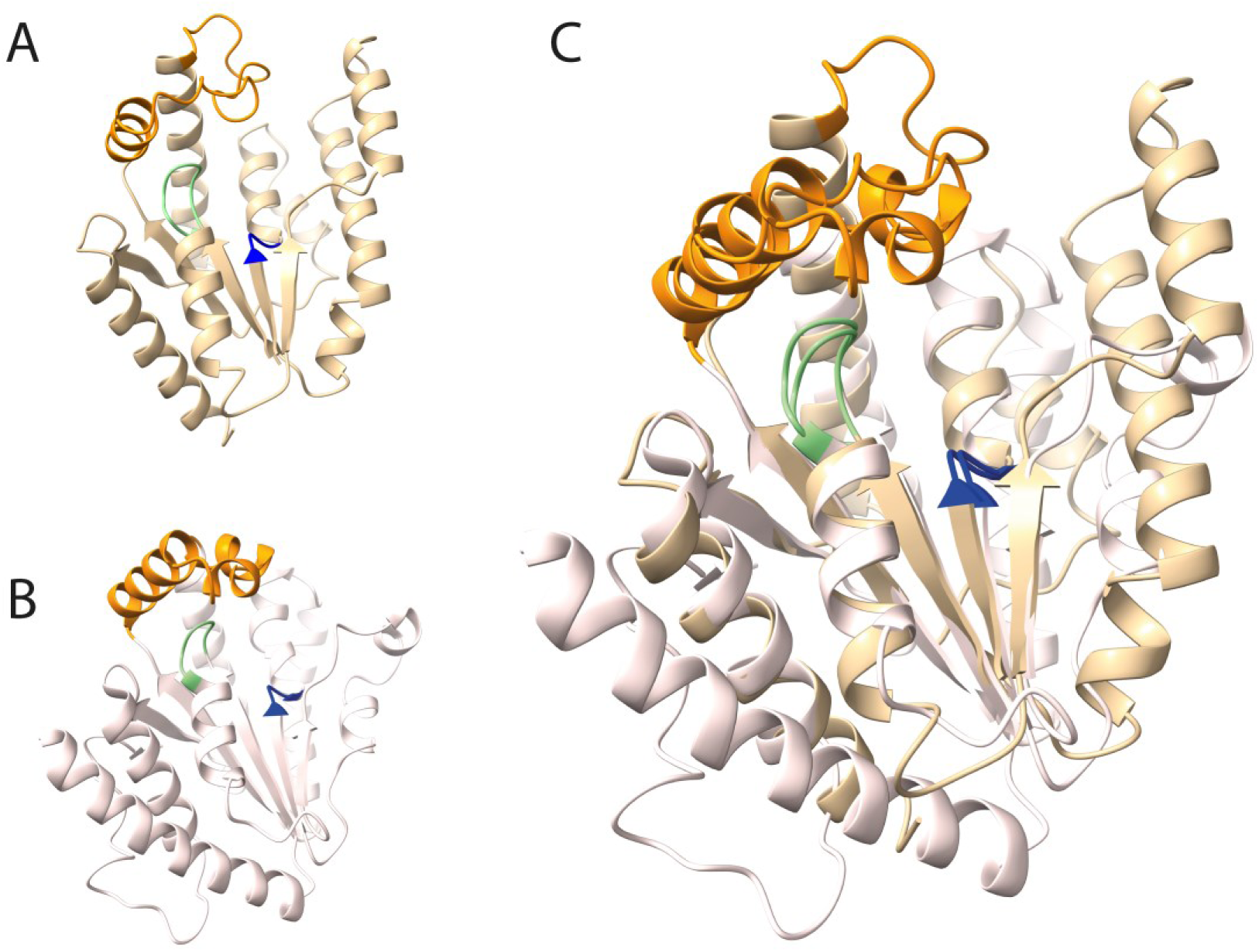
Structural comparison of the *S. acidocaldarius* cPPK3 (Saci_2019) with PPK2 from *Francisella tularensis*. The ribbon representations of the monomers of Saci_2019 (alphafold model, brown) (A) and the *F. tularensis* PPK2 (crystal structure, pdb 5llb, light pink) (Parnell, Mordhorst et al. 2018) (B) are shown. The superimposition (C) clearly shows the structural differences in the N-terminal α helix and the distal helix bundle between cPPK3 from *S. acidocaldarius* and the PPK2s as well as their similarity in the core fold (the structural motifs characteristic to the P-loop NTPases, i.e. the Lid region, the P-loop and the DRX motif (Leipe, Koonin et al. 2003), are highlighted in orange, green and blue, respectively).

As revealed by BLAST searches, homologues of PPK3 and rPPK3s have a narrow distribution and are mainly restricted to Sulfolobales and some Nitrososphaerales, both from the archaeal TACK group. Only few bacterial homologues were identified among others many of “Candidatus” organisms or from members of the Acidobacteriota. The PPK3s and rPPK3s share 22-38% sequence identity to each other and 10-30% sequence identity to canonical dTMPKs. To PPK2s sequence identities in the range of only <18% were found. In accordance with this, phylogenetic analyses (Fig. 6) clearly revealed a closer relationship of (r)PPK3s with canonical dTMPKs forming a monophyletic group clearly separated from PPK2s. However, cPPK3s and rPPK3s form two separate subgroups more closely related to each other than to the dTMPKs suggesting that both groups might have originated from dTMPKs via two gene duplication events in the TACK group. As previously observed (Motomura, Hirota et al. 2014) the PPK2s cluster according to the three described classes with rather deep branching class II enzymes forming a badly resolved group between the clearly separated class I and class II PPK2s also previously observed (Leipe, Koonin et al. 2003, Motomura, Hirota et al. 2014).

**Figure 6.**
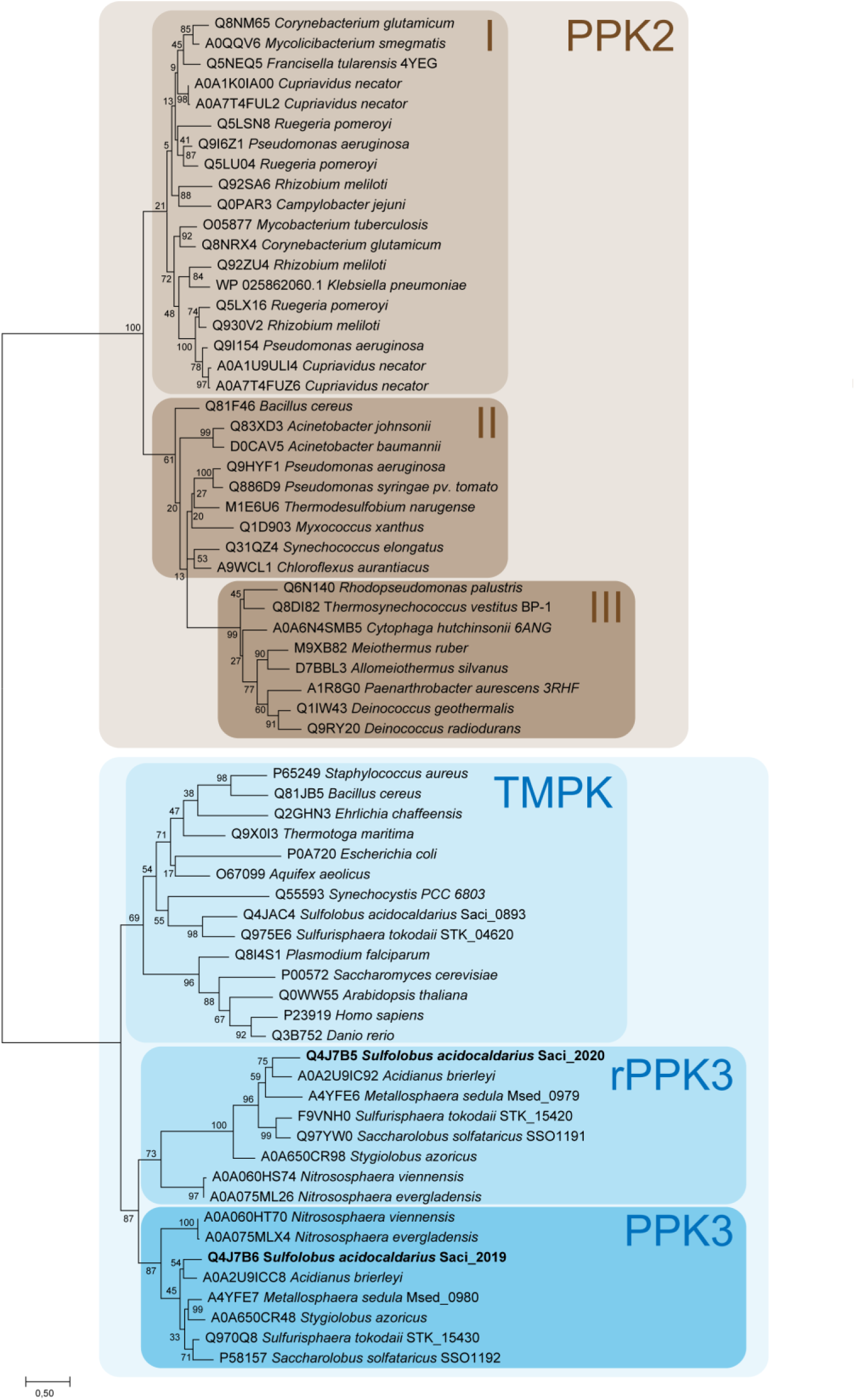
Phylogenetic affiliation of PPK3s with dTMPKs and PPK2s. Organism names and uniport accession numbers are given and enzymes characterized in this study are marked in bold-face. Class I, II and III PPK2s (Neville, Roberge et al. 2022) are highlighted in light brown, brown, and dark brown, respectively. The group of dTMPK like proteins harbors the canonical dTMPKs (light blue), and the PPK3s which are subdivided into the regulatory, inactive rPPK3s (blue) and the catalytic active cPPK3s (dark blue). The evolutionary history was inferred by using the Maximum Likelihood method and Le_Gascuel_2008 model (Le and Gascuel 2008). The model was selected based on the lowest Bayesian information criterion value using the “Find best protein model” option implemented in the MEGA11 package. The tree with the highest log likelihood (−13446,63) is shown. The percentage of trees in which the associated taxa clustered together is shown next to the branches. Initial tree(s) for the heuristic search were obtained automatically by applying Neighbor-Join and BioNJ algorithms to a matrix of pairwise distances estimated using the JTT model, and then selecting the topology with superior log likelihood value. A discrete Gamma distribution was used to model evolutionary rate differences among sites (4 categories (+*G*, parameter = 1,6717)). The tree is drawn to scale, with branch lengths measured in the number of substitutions per site. This analysis involved 66 amino acid sequences. All positions containing gaps and missing data were eliminated (complete deletion option). There were a total of 148 positions in the final dataset. Evolutionary analyses were conducted in MEGA11 (Tamura, Stecher et al. 2021).

Sequence and structural comparisons revealed that in (r/c)PPK3s as shown in the crystal structure of the *S. tokodaii* STK_15430 PPK3, ATP/ADP binding occurs in the similar position as in dTMPKs (e.g. *E. coli* and *Aquifex aeolicus*) (Ostermann, Schlichting et al. 2000, Biswas, Shukla et al. 2017, Biswas, Shukla et al. 2017) and the binding residues are highly conserved (see Alignment in Figure W and structural comparison in SI Figure 14). Conversely, the residues involved in TMP binding including the EP motif (Biswas, Shukla et al. 2017) in dTMPKs are not conserved in (r/c)PPK3s which appears plausible with regard to the different second substrate (TMP in TMPKs vs. polyP in PPK3s). Instead, structural analyses identified a positively charged cleft in PPK3s (Fig. 7 and SI Fig. 15), which could accommodate polyphosphate as a ligand as revealed by docking experiments. The residues forming this positively charged cleft and likely contributing to polyP binding (shown in SI Figure 14) are highly conserved in PPK3 sequences but not in rPPK3s. This extended positively charged binding cleft is analogous to PPK2s (E) although the binding sites for ADP and polyP are switched. Also, the catalytically essential arginine residue from the DRX motif functioning as clamp in binding and correct positioning of the terminal phosphate moieties from donor and acceptor in TMPKs and PPK2s (Ostermann, Schlichting et al. 2000, Biswas, Shukla et al. 2017, Neville, Roberge et al. 2022) is absent in rPPK3s. These findings explain the missing enzymatic activity of the Saci_2020 rPPK3.

**Figure 7.**
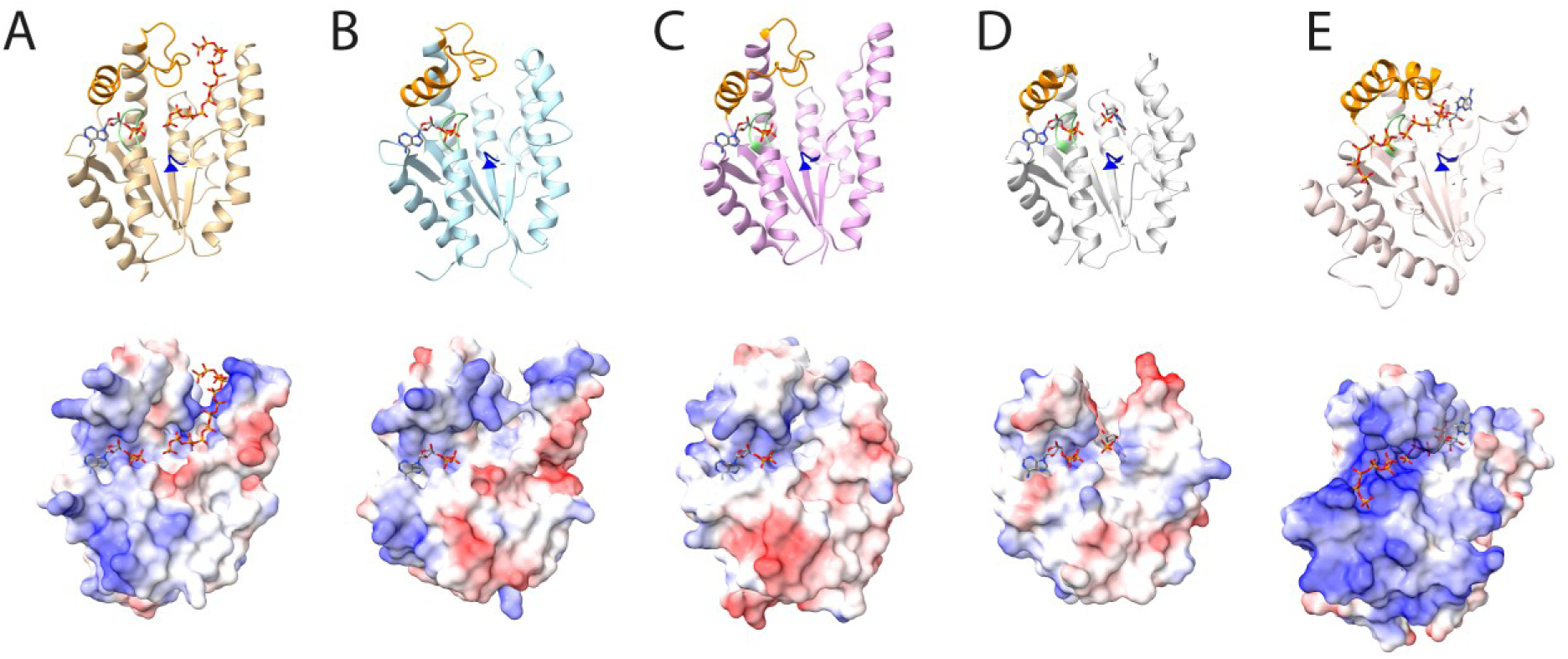
Substrate binding sites in PPK3s in comparison to *A. aeolicus* dTMPK and *F. tularensis* PPK2. Upper panels show the ribbon representation, and lower panels the electrostatic surface representation (blue positive charge, red negative charge) of (A) Saci_2019 cPPK3 (alphafold model), (B) Saci_2020 rPPK3 (alphafold model), (C) *S. tokodaii* cPPK3 (STK_15430) (crystal structure, pdb 5h70) (Biswas, Shukla et al. 2017), (D) *A. aeolicus* dTMPK (crystal structure, pdb 5xb2) (Biswas, Shukla et al. 2017), and the *F. tularensis* PPK2 (crystal structure, 5llb) (Parnell, Mordhorst et al. 2018). ADP, TMP (D), as well as polyP_6_ and ADP substrate analogue (β,γ-methylene adenosine 5ʹ-pentaphosphate) (E) are shown as stick models. In Saci_2019 and Saci_2020 the ADP is shown in the same position as in the crystal structure of STK_15430 from *S. tokodaii* (Biswas, Shukla et al. 2017). The proposed polyphosphate binding site in the Saci_2019 cPPK3 (A) (polyphosphate shown as stick model) was identified by swissdock (Grosdidier, Zoete et al. 2011, Bugnon, Röhrig et al. 2024) based docking experiments using the polyP_9_ available in the pdb database (9PI) as ligand. Two of the best hits are shown as stick models (with overlapping phosphate moieties after manual adjustments). In the PPK3s Saci_2019 and STK_15430 polyphosphate binding is proposed to take place in a positively charged binding cleft (upper right of the structures) which is missing in the rPPK3 Saci_2020 further explaining the missing enzymatic activity. Also, in the dTMPK from *A. aeolicus* this positively charge region is missing instead showing a more hydrophobic binding pocket for the thymidine moiety of TMP. This extended positively charged binding cleft is analogous to PPK2s (E) although the binding sites for ADP and polyP are switched. A detailed close-up view of the proposed ADP and polyP binding site for Saci_2019 PPK3 is shown in SI Fig. 15.

Size exclusion chromatography indicated an α_2_β_2_ heterotetrameric structure of *Sa*PPK3. The alphafold model (Abramson, Adler et al. 2024) suggest that a rPPK3 Saci_2020 dimer makes up the core of the complex to which one Saci_2019 cPPK3 is attached to both sides (Fig. 8). Notably, AlphaFold 3 correctly predicts the position of ADP which adopts the same position as observed in the crystal structure of STK_15430 supporting the model prediction (Biswas, Shukla et al. 2017). Interestingly, the Lid domain is predicted to be involved in the interaction of both Saci_2020 subunits. Furthermore, each of the Saci_2019 subunits interact with both subunits of the Saci_2020 dimer. This is in contrast to canonical dTMPKs which usually form homodimers. Nevertheless, Saci_2019 and Saci_2020 have a similar spatial orientation to each other as both subunits of the homodimer in TMPKs, however with subtle differences. It appears reasonable to speculate that the tetramer formation might enable the Saci_2019 activation by Saci_2020 by favoring/stabilizing the active conformation through structural interaction.

**Figure 8.**
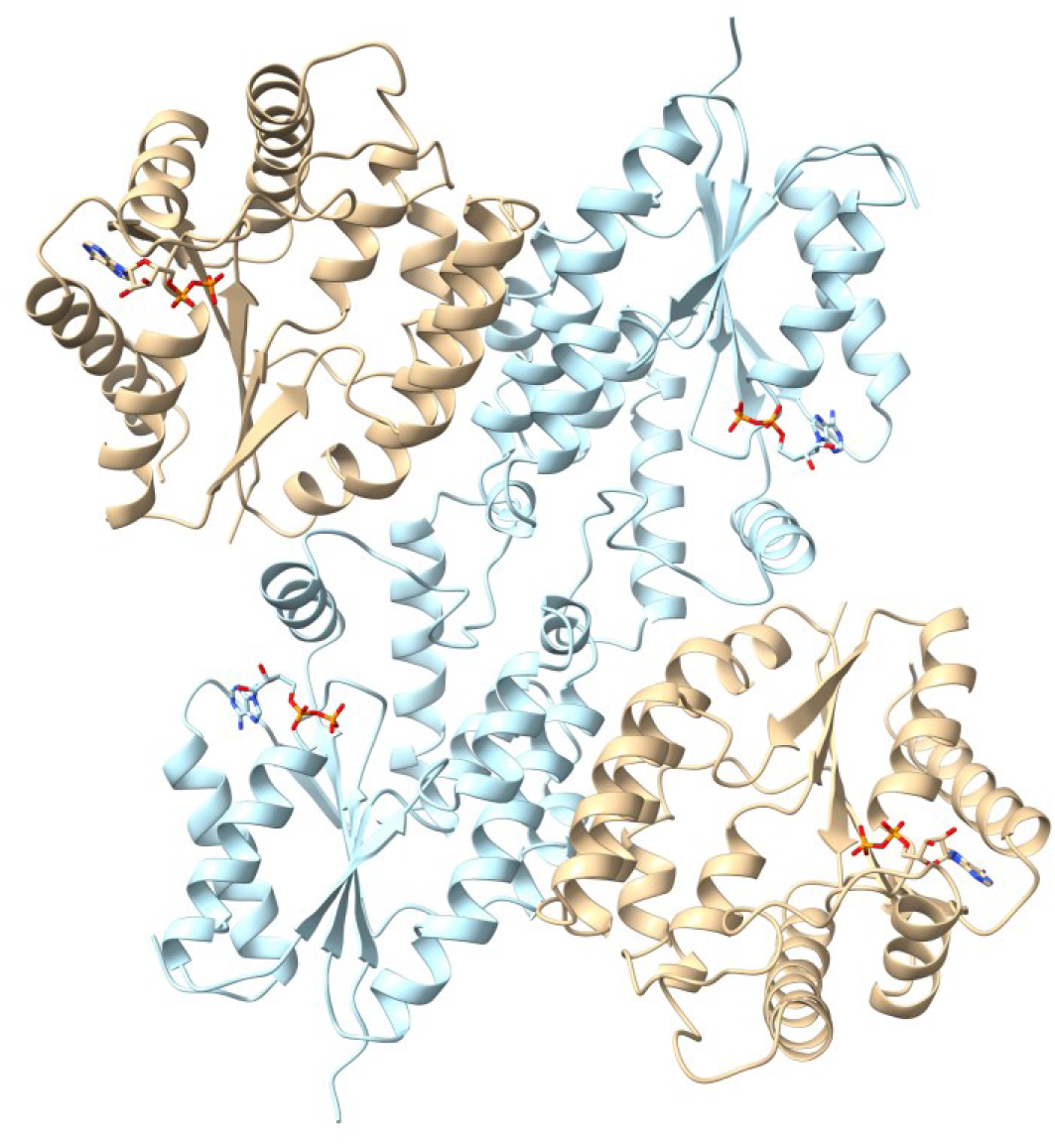
Structural model of the heterotetrameric *Sa*PPK3. The protein is composed of two of each subunits Saci_2019 cPPK3 (brown) and Saci_2020 rPPK3 (blue) and the oligomeric structure was revealed by size exclusion chromatography. The model was generated using the alphafold server (Abramson, Adler et al. 2024) with ADP as a ligand. The model suggests that heteromer formation is enabled by the Saci_2020 subunit.

## Discussion

PolyP has been found in all three domains of life, however, although its function as phosphate and energy storage as well as regulator of a multitude of cellular processes is well established in Bacteria and Eukaryotes the knowledge of polyP metabolism in Archaea is only limited. The presence of polyP has been demonstrated in Archaea and the effect of low polyP levels (*ppx* overexpression) on motility, adhesion, biofilm formation and heavy metal resistance has been proven in *Sulfolobus spp.* (Recalde, van Wolferen et al. 2021). Contrary to this functional analysis, however, the enzymes involved in polyP metabolism have hardly been studied so far. Bioinformatic studies provide an incomplete picture of enzymes involved in polyP metabolism in many archaea (Orell, Navarro et al. 2012, Paula, Chin et al. 2019, Wang, Liu et al. 2019). In all members of the Crenarchaeota homologues for known PPK of the PPK1 and PPK2 and Arps are missing, suggesting the presence of novel enzymes.

The genome neighborhood of the gene encoding the reported PPX (*saci_2018)* revealed a conserved gene organization resembling the one in Bacteria and other Archaea, although classical PPK1 and/or PPK2 enzymes were missing. Available transcriptome data confirmed that the *ppx* gene forms an operon with genes encoding SixA (*saci_2017*) two thymidylate kinases (dTMPKs, *saci_2019* and *saci_2020*) as well as a CHAD-domain containing protein (*saci_2021*). SixA is a member of the phosphoglycerate mutase histidine phosphatase superfamily and for CHAD domain proteins a function as polyP binding module that assists in substrate recruitment and enhances the processivity of polyP-metabolizing enzymes and/or a role in copper resistance via divalent metal ion binding has been proposed (Tumlirsch and Jendrossek 2017, Lorenzo-Orts, Hohmann et al. 2019, González-Madrid, Navarro et al. 2024), however, their role in Archaea remains to be elucidated. A third dTMPK-encoding gene was identified at a different locus in the *S. acidocaldarius genome* (*saci_0839*).

Since dTMPKs have been characterized as the closest structural homologs of PPK2 (Nocek, Kochinyan et al. 2008, Motomura, Hirota et al. 2014) we expressed and analyzed the three recombinant *Sa*dTMPKs for their enzymatic activity. Of the three *Sa*dTMPKs only Saci_0893 possessed thymidylate kinase activity, whereas cPPK3 Saci_2019 revealed PPK activity that was significantly enhanced by the regulatory subunit rPPK3 Saci_2020, which showed neither dTMPK or PPK activity by itself. Titration and gel filtration experiments revealed an optimal ratio of 1:1 for Saci_2019 and Saci_2020 proteins and complex formation with a tetrameric structure of *Sa*PPK3 (α_2_β_2_), which was supported by model predictions using alphafold (Fig. 7). For cPPK3 Saci_2019 alone a monomeric structure was observed. The enzyme required divalent metal enzymes (i.e. Mg^2+^) for activity. *Sa*PPK3 activity was shown to be reversible with clear preference (50-fold higher specific activity, Table 1 (Jacky)) for polyP-dependent nucleotide kinase activity, i.e. polyP-dependent formation of ATP from ADP. In both directions, polyP and nucleotide synthesis the activity of cPPK3 Saci_2019 was significantly enhanced by addition of the regulatory subunit rPPK3 Saci_2020. In addition to ADP also GDP was used as P_i_ acceptor forming GTP, however, with significantly reduced activity (x-fold), whereas no activity in the direction of polyP synthesis was observed using GTP as Pi donor.

To unravel the kinetic and mechanistic properties of *Sa*PPK3 we used a combined experimental and modelling approach (Shen, Kohlhaas et al. 2020, Kuschmierz, Shen et al. 2022) with quantitative ^31^P-NMR. For model construction, we determined the initial enzyme rate kinetics of the reversible *Sa*PPK3 by enzyme characterization and metabolite (i.e. polyP and PEP) stability to identify additional sources of Pi release via ^31^P-NMR. For polyP the non-enzymatic formation of P_i_ and cP_3_ from polyP in absence of enzymes and presence of metal ions and for PEP an enhanced P_i_ release in presence of *Sa*PPK3 was observed that was included into our model.

The non-enzymatic formation of cyclic polyPs (cP_3_-P_6_) at high temperature and in presence of metal ions has been reported previously (Rulliere, Perenes et al. 2012) and more recently also the presence of cyclic polyPs *in vivo* has been shown (Mandala, Loh et al. 2020). Although their physiological role is still unknown it is tempting to speculate that they possess a role as signal molecules/second messengers (Blank 2023). So far in *Saci* only few nucleotide-based second messengers such as 3’,5’-cyclic adenosine and guanosine monophosphate (cAMP, cGMP) and diadenosine tetraphosphate (Ap4A) were identified whereas typical bacterial second messengers cyclic diguanosine monophosphate (3’,5’-c-di-GMP) and guanosine (penta-)/tetraphosphate ((p)ppGpp) are absent, however, their regulatory function still needs to be elucidated (van der Does, Braun et al. 2023).

In line with the preferred polyP-dependent nucleotide kinase activity *Sa*PPK3 catalyzes the polyP-dependent ATP formation at high velocity whereas polyP formation is only observed in presence of an energy recycling system.

Notably in all *Sa*PPK3 assays (both directions) an ATP/ADP equilibrium ratio of 3-4 is observed, which has been also reported in a similar range (2.3) for bacterial PPK1s from *E. coli* and *Vibrio cholerae* and PPK2 from *Sinorhizobium meliloti (renamed Ensifer meliloti*) in a biocatalytic study on ATP recycling systems (Keppler, Moser et al. 2022). Unfortunately, no information is available for *Saci*, but in respiring *E. coli* cells an ATP/ADP ratio of ∼30 (∼3.5 mM ATP and 0.12 mM ADP) was determined (Meyrat and von Ballmoos 2019). Hence, our studies suggest a function of polyP as energy buffer mediated by PPKs only under low energy charge of the cell rather than an active role in buffering ATP/ADP during active metabolism. This seems to be in line with the observation that PPKs are not essential and physiological studies using *ppk* deletion strains show diverse phenotypes often related to stress response or energy-dependent cellular processes such as motility and biofilm formation (Rao, Gómez-García et al. 2009). For example, in *E. coli* stationary phase cells the deletion of the *ppk1* gene resulted among others in decreased long-term survival and increased sensitivity to heat, oxidative and osmotic stress (Rao and Kornberg 1996, Xie and Jakob 2019). In addition, a close relationship of polyP metabolism to multiple regulatory pathways such as nitrogen and P_i_ limitation as well as stringent response was reported (Ault-Riché, Fraley et al. 1998, Rao and Kornberg 1999). In *R. eutropha* all seven PPKs could be deleted and no phenotype in response to heat stress, bleach nor an effect on motility or biofilm formation was observed in complex medium (Rosigkeit, Kneißle et al. 2021). However, due to the multiple functions of polyP as P_i_ store, metal ion chelator and chaperone and pro-aggregation molecule it remains difficult to attribute cellular responses to a certain function.

Sequence, structural and phylogenetic analysis reveal that *Sa*PPK3 (α_2_β_2_) homologues form a novel PPK3 family in the thymidylate kinase (dTMPK) family and are only distantly related to PPK2s both within the P-loop kinase superfamily and not related at all to PPK1 (Leipe, Koonin et al. 2003). Both PPK2s and PPK3s possess the typical structural features of P-loop kinases i.e. the Lid region, the P-loop (Walker A motif) and the DRX (Walker B) motif, however, as shown here, there are significant sequence and structural differences. Notably the structure of the *S. tokodaii* homologue of the cPPK3 subunit 2019 (STK_15430), which is part of the same polyP operon as in *S. acidocaldarius*, has been solved, and the enzyme was reported to function as unusual thymidylate kinase (Biswas, Shukla et al. 2017). However, the authors monitored only ADP formation from ATP and not dTDP formation, overlooking both the PPK activity and the presence of two other dTMPKs in *S. tokodaii*. The composition with a catalytic active and regulatory inactive subunit of *Sa*PPK3 resembles PPK2 class II two-domain fusion proteins (e.g. PPK2C *P. aeruginosa* (PA3455), PSPTO1640 from *P. syringae* pv. tomato and PPK2 *Acinetobacter johnsonii*) with double size (Shiba, Itoh et al. 2005, Nocek, Kochinyan et al. 2008). The *P. aeruginosa* class II PPK2C was shown to have a catalytically inactive N-terminal and active C-terminal domain for which a function in protein dimerization (four domain structure) has been proposed (Nocek, Kochinyan et al. 2008). Similar to our results for the inactive rPPK3 domain, the DRX (Walker B) motif is absent in the N-terminal domain and a different distribution of positively charged residues on protein surface was observed. Notably for all two domain PPK2 class IIs tested only polyP-dependent AMP kinase activity catalyzing the polyP-dependent phosphorylation of AMP (GMP) to ADP (GDP) was reported, whereas one domain PPK2 (e.g. SMc02148 from *Sinorhizobium meliloti*) catalyze the polyP-dependent phosphorylation of ADP to ATP. The reverse reaction in the direction of polyP synthesis (in absence of a, energy recycling system) was not observed (Shiba, Itoh et al. 2005, Nocek, Kochinyan et al. 2008). Hence despite the different structure and enzyme activity with either ADP (*Sa*PPK3) or AMP (2 domain PPK2 class IIs) both PPKs seem to utilize the regulatory subunit for protein dimerization and thus activation.

In summary, compared to bacteria and eukaryotes, relatively little is known about polyP metabolism and functions in Archaea. In *S. metallicus* and *S. solfataricus*, the influence of polyP accumulation in response to metal toxicity (e.g. copper) was studied due to its role in bioleaching (Remonsellez, Orell et al. 2006, Rivero, Torres-Paris et al. 2018, Soto, Recalde et al. 2019). Furthermore, in *S. acidocaldarius*, a reduction in polyP by *Sa*PPX overexpression has been linked to decreased motility, adhesion and biofilm formation (Recalde, van Wolferen et al. 2021). However, a detailed characterization of the PPX is still missing. In addition, a highly active triphosphatase (triphosphate tunnel metalloenzyme (TTM), Saci_0718) that uses polyP_3_ and polyP_4_ as preferred substrates and converts them to PP_i_ and P_i_ (Vogt, Ngouoko Nguepbeu et al. 2021) as well as a cytoplasmic inorganic pyrophosphatase forming Pi from PPi (Saci_0955, family I pyrophosphatase) (Meyer, Moll et al. 1995) have been reported from *S. acidocaldarius*. Therefore, future work is required to elucidate the polyP metabolism in Archaea. In addition, the dual function of polyP as P_i_ storage or energy buffer requires a sophisticated synthesis/degradation regulation in order to adopt to fluctuations in the environment and internal demands closely linked to P_i_ and energy homeostasis, which still has to be elucidated in archaea. Finally, the identification of the novel PPK3 family in members of the *Sulfolobales* highlights once again the endowment of archaea with novel enzymes from different enzyme families that perform the same function as their bacterial or eukaryotic counterparts.

## Materials and methods

### Strains and growth conditions

The *Escherichia coli* strains DH5α for cloning were grown in Luria-Bertani (LB) medium, whereas Rosetta (DE3) strains were grown in Terrific broth (TB) medium (22 g/L yeast extract, 12 g/L tryptone/peptone from casein, 0.4 % (v/v) glycerol), containing the appropriate antibiotics (100 µg/mL ampicillin (pET15b), 50 µg/mL kanamycin (pET28b), 25 µg/mL chloramphenicol (Rosetta (DE3))) at 37 °C.

### Cloning, heterologous expression and purification of the recombinant proteins

The *ppx* (*saci*_2018, GenBank: AAY81314.1), *dtmpk* (saci_0893, GenBank: AAY80256, *saci*_2019, GenBank: AAY81315.1, and *saci*_2020, GenBank: AAY81316.1) genes were amplified via PCR using genomic DNA of *S. acidocaldarius* DSM639 as template (for primers see SI Tab. 1). PCR fragments were cloned into the vector pET15b (*ppx*, *dtmpk*:saci_0893, saci_2019) and pET28b (*dtmpk*:saci_2020) and successful cloning was confirmed by sequencing. The genes were cloned with N-terminal 6x His-Tag which originates from the pET vector system. The expression vectors were used for heterologous gene expression in *E. coli* Rosetta (DE3) cells grown on TB-medium and the induction was performed at OD_600_ 0.5 - 0.6 with 1 mM isopropyl-β-d-thiogalactopyranoside (IPTG, Carl Roth GmbH + Co. KG, Karlsruhe, Germany). After incubation overnight at 18 °C with shaking (180 rpm) and cell harvest (6.000 x *g*, 15 min, 4 °C), the cell pellet (1 g cells (wet weight)) was resuspended in 3 ml 50 mM TRIS/HCl, 300 mM NaCl, pH 8.0 supplemented with protease inhibitor (Merck KGaA, Darmstadt, Germany) and disrupted by French press (20.000 psi). In addition, thermally unstable *E. coli* proteins were removed by heat precipitation (20 min at 70 °C with gentle agitation) followed by centrifugation (25.000 x *g*, 45 min, 4 °C). The cleared lysate with the His-tagged proteins were purified via Protino^®^ Ni-TED column (Machery & Nagel, Düren, Germany) following the instructions of the manufacturer. 10 mM TRIS/HCl, 300 mM NaCl, 250 mM imidazole, pH 8.0 was used as elution buffer. Protein concentrations were determined with the standard Bradford assay by measuring the absorption with microplate reader (Infinite M200, TECAN) at 450 and 595 nm (Bradford 1976, Zor, Selinger 1996). The pure proteins were stored in 10 mM TRIS/HCl, 300 mM NaCl, 250 mM imidazole, 15 % (v/v) glycerol, pH 8.0 at - 80 °C and used for further enzymatic characterization. Purification and molecular mass of subunits were monitored by SDS-polyacrylamide gel electrophoresis and Coomassie Brilliant Blue staining.

To determine the native molecular mass of the recombinant proteins, a calibration curve was generated by running ribonuclease A (13.7 kDa), ovalbumin (44 kDa), conalbumin (75 kDa), aldolase (158 kDa) and ferritin (440 kDa) from the Gel Filtration LMW and HMW Calibration Kit (Cytavia, USA). The size exclusion chromatography was performed for cPPK3 (Saci_2019) and *Sa*PPK3 (Saci_2019 and Saci_2020 (1:1 ratio)) (Superose® 6 Increase 10/300 GL, Cytiva Lifescience™, Freiburg, Germany) using 50 mM TRIS/HCl with 300 mM NaCl, pH 8.0 as buffer. The oligomeric state of native cPPK3 (Saci_2019) and SaPPK3 (Saci_2019 and Saci_2020 was calculated using the generated calibration curve resulting in a molecular weight of 21.3 kDa for cPPK3 (Saci_2019) and 114.2 and 23.8 kDa for SaPPK3 (Saci_2019 and Saci_2020).

### ^31^P NMR for analysis of predicted dTMPK activity

For qualitative analysis via ^31^P NMR the reaction mixtures contained 2 mM ATP, 1 mM dTMP, 4 mM MgCl_2_, 50 mM KCl, 10 µg of the respective enzyme (Saci_0893, Saci_2019 or Saci_2020, respectively) in 50 mM TRIS/HCl pH, 7.4 with 10 % (v/v) D_2_O in a total volume of 1 ml. The samples were incubated at 70 °C for 8 hours and analyzed via ^31^P NMR spectroscopy.

### Enzyme assays following ADP, P_i_ and polyP formation

To further analyze the enzymatic activity of Saci_0893, Saci_2019, Saci_2020, or Saci_2019 and Saci_2020 (1:1)) discontinuous assays were performed following the formation of ADP, P_i_ and polyP using the pyruvate kinase (PK) – L-lactate dehydrogenase (LDH) assay, malachite green assay (for details see below) and polyP assay (MicroMolar Polyphosphate Assay Kit, ProFoldin), respectively. Therefore, the dTMPK enzyme assay was performed at 70°C with 1 mM dTMP, 2 mM ATP, 50 mM KCl, 4 mM MgCl_2_ and 10 µg/ml Saci_0893 in 50 mM TRIS/HCl, pH 7.4 at 70 °C in a total volume of 1 ml (Biswas, Shukla et al. 2017). Aliquots (120 µl) were removed at 0, 15, 30, 45 and 60 min and the indicator reactions were performed at 37°C.

#### PK-LDH assay to follow ADP formation (polyP or dTDP formation)

The enzyme assay couples the formation of ADP to NADH oxidation via PK (rabbit muscle, (Merck, Darmstadt, Germany)) and LDH (rabbit muscle, (Merck, Darmstadt, Germany)). The indicator reaction was performed at 37°C for 15 min with 50 µL of sample volume, 2 mM phosphoenolpyruvate (PEP), 4 mM MgCl_2_, 50 mM KCl, 0.5 mM NADH, 8 U/ml PK, and 4 U/ml LDH in 50 mM TRIS/HCl pH 7.4 in 200 µl total volume. The NADH consumption was detected at 340 nm using a microplate reader (Infinite M200, TECAN), with an internal calibration curve generated for 0 - 2 mM ADP. One unit (1U) of enzyme activity is defined as 1 µmol of product (ADP) formed per min.

#### Malachite green assay to follow Pi formation

Malachite green reagent was prepared using 1.5 mL of 1 % (w/v) malachite green oxalate, 30 mL of 1 % (w/v) ammonium molybdate both solved in distilled H_2_O, 4.64 mL of 70 % (v/v) HClO_4_, 13.86 mL of H_2_O. After stirring 30 min the solution was supplemented with 0.45 mL of 5 % (v/v) Triton X-450 solved in H_2_O and filtered (Andreeva, Ledova et al. 2019). The malachite green reagent was stable at room temperature for up to 6 months. The malachite green assay was performed by adding 20 µL of sample volume to 300 µL of the malachite green reagent and following incubation at room temperature for 20 min. The absorption was determined at 650 nm using microplate reader (Infinite M200, TECAN). A standard curve was prepared using 0-500 µM KH_2_PO_4_ (following (Trilisenko, Andreeva et al. 2015)) using 20 µL sample volume and 300 µL malachite green reagent. The P_i_ concentrations were calculated with the generated standard curve.

#### PolyP assay to follow polyP formation

The MicroMolar Polyphosphate Assay Kit (ProFoldin, Massachusetts, United States) was used for determination of the polyP concentration according to the manufacturer’s instructions. Briefly, 30 µL of samples were mixed with 30 µL of the fluorescent dye (PPD dye, prepared following manufacturer’s instructions) and fluorescence intensity was determined using a microplate reader (Infinite M200, TECAN) (emission 550 nm, excitation 415 nm) using 0 - 50 µM polyP_45_ as standard. One unit (1 U) of enzyme activity is defined as one µmol of product (polyP) formed per min.

#### Polyphosphate kinase (PPK) *Enzyme assays for the forward reaction of polyP formation*

PPK activity was determined in a discontinuous assay as described previously (Keasling and Hupf 1996). Briefly, the enzyme reaction was performed at 70°C in 50 mM TRIS/HCl, pH 7.4 with 50 mM KCl and 4 mM MgCl_2_ in the presence of 20 µg/ml enzyme (Saci_2019, Saci_2020, or Saci_2019 and Saci_2020 (1:1)) in a total volume of 1 ml. After preincubation for 90 seconds at 70 °C, the reaction was started by addition of the substrate ATP (2 mM) for polyP synthesis. During the reaction, aliquots were taken every 15 min over a time period of 90 min and the produced ADP, polyP and P_i_ were determined in the indicator reaction at 37°C or RT using the discontinuous PK-LDH assay and the malachite green assay, respectively (see above for details).

To determine the optimal molar ratio of both proteins (Saci_2019 and Saci_2020) the PPK3 assay (for polyP synthesis) following ADP formation was performed in presence of 10 µg/ml Saci_2019 and 0 – 10 µg/ml Saci_2020 were added in a total reaction volume of 0.5 ml at 70°C. Additionally, a titration experiment was performed using both enzymes in different ratios from 10:0 to 0:10 (20 µg/ml total protein). The indicator reactions were carried out at 37°C or RT to detect the formation of polyP (polyP assay) and ADP (PK-LDH assay) as described above.

The temperature dependence of the heteromeric *Sa*PPK3 (10 µg/ml Saci_2019 and 10 µg/ml Saci_2020 (1:1)) was determined in the range of 50 to 90 °C at pH 7.4. The NADH consumption was detected at 340 nm using a microplate reader (Infinite M200, TECAN), with an internal calibration curve generated for 0 - 2 mM ADP. One unit (1U) of enzyme activity is defined as 1 µmol of product (ADP) formed per min (adjusted at the respective temperatures). The pH dependence was analyzed for heteromeric *Sa*PPK3 in a mixed buffer system (50 mM TRIS/HCl, Citrate/HCl, BICINE/NaOH and MES/NaOH) in the pH range of 4 – 9 at 70 °C.

The kinetic characterization of *Sa*PPK3 in the direction of polyP formation was performed in presence of 0 - 8 mM ATP (homomeric cPPK3, Saci_2019) and 0 - 4 mM ATP (heteromeric *Sa*PPK3, Saci_2019 and Saci_2020 (1:1)) with 4 mM MgCl_2_, 50 mM KCl and 10 µg/ml of enzyme in 50 mM Tris/HCl, pH 7.4 at 70 °C in a total volume of 0.5 ml. Aliquots (50 µl) were removed every 15 min over a period of 90 min. ADP formation was determined via the discontinuous PK-LDH assay (see above). Enzyme activity with GTP was tested using 2 mM GTP instead of ATP as substrate. To investigate ADP inhibition 2.5 mM ADP were added to the PPK assay containing 1.5 mM ATP as substrate and 10 µg/ml heteromeric *Sa*PPK3 (0.5 ml total volume) and polyP formation was determined using the discontinuous polyP assay (see above).

#### Polyphosphate kinase (PPK) Enzyme assays for the **reverse reaction** of nucleotide formation

Reverse PPK activity was assayed using a discontinuous assay following ATP or GTP formation and polyP degradation. The assay was performed at 70°C in presence of 2 mM ADP or GDP, 50 µM polyP_45_, 10 mM MgCl_2_, 50 mM KCl and 5 µg/ml of homomeric cPPK3 (Saci_2019) or heteromeric *Sa*PPK3 (Saci_2019 and Saci_2020 (1:1)) in 50 mM TRIS/HCl, pH 7.4 and a total volume of 1 ml. The assay was performed for 90 min and aliquots (100 µl) were removed every 15 min. The indicator reaction was performed at 37°C or RT and the formed ATP and polyP was determined via the HK-G6PDH assay (see below) and polyP assay (see above), respectively.

HK-G6PDH assay to follow NTP formation. PPK3 activity for NTP formation was determined by coupling the ATP/GTP formation to the NADP^+^ reduction via HK (yeast, (Roche, Basel, Switzerland)) and G6PDH (yeast, (Roche, Basel, Switzerland))). The indicator reaction was performed at 37°C for 15 min using 50 µL sample volume, 2 mM glucose, 10 mM MgCl_2_, 50 mM KCl, 500 µM NADP^+^, 1 U/ml HK and 1 U/ml G6PDH in 50 mM TRIS/HCl, pH 7.4 in a total volume of 200 µL. The NADPH formation was detected at 340 nm with an internal calibration curve for (0 – 2 mM) ATP or GTP, using a microplate reader (Infinite M200, TECAN). One unit (1 U) of enzyme activity is defined as 1 µmol of product (ATP or GTP) formed per min.

The kinetic characterization in the direction of nucleotide formation was performed via the discontinuous HK-G6PDH assay in presence of 0-40 mM ADP with 100 µM polyP_45_ (for ADP) and 0-100 µM polyP_45_ with 20 mM ADP (for polyP) in presence of 10 mM MgCl_2_, 50 mM KCl and 10 µg/ml of heteromeric *Sa*PPK3 (Saci_2019 and Saci_2020 (1:1)) *Sa*PPK3 in 50 mM TRIS/HCl, pH 7.4 in a total volume of 0.5 ml.

#### Thyidilate kinase (dTMPK)

For characterization of the dTMPK Saci_0893 a continuous dTMPK assay was performed with varying concentrations of dTMP (0.05 – 1 mM), 2 mM ATP, 4 mM MgCl_2_ (optimal ATP to MgCl_2_ ratio), 0.2 mM NADH, 2 mM PEP, 8 U PK (rabbit muscle, (Merck, Darmstadt, Germany)), 4 U LDH (rabbit muscle, (Merck, Darmstadt, Germany)) and 1.25 µg/ml of Saci_0893 in 50 mM TRIS/HCl, pH 7.4 at 55°C in a total volume of 500 µl. The NADH consumption was detected spectrophotometrically using a Specord UV/visible-light (Vis) spectrometer (Analytic Jena, Germany) at 340 nm (extinction coefficient of NADH 5.8 mM^-1^ cm^-1^ at 37°C). One unit was defined as 1 µmol NADH oxidized per min.

### Analysis of the reversible heteromeric *Sa*PPK3 using ^31^P NMR spectroscopy

For qualitative analysis, the PPK enzyme reactions and control samples were followed via ^31^P NMR measurements at 70 °C for up to 12 hours. The assay following ATP formation was performed in presence of 10 mM ADP, 500 µM polyP_45_, 10 mM MgCl_2_, 50 mM KCl with 20 µg/ml of heteromeric PPK (10 µg/ml Saci_2019 and 10 µg/ml Saci_2020, respectively) in 50 mM TRIS/HCl, pH 7.4 with 10 % (v/v) D_2_O (1 ml total volume, 750 min). The reaction mixture following polyP synthesis was performed in presence of 2 mM ATP, 4 mM MgCl_2_, 50 mM KCl, with 20 µg/ml of heteromeric *Sa*PPK3 in 50 mM TRIS/HCl, pH 7.4 with 10 % (v/v) D_2_O (1 ml total volume, 360 min). The PPK assay for polyP formation with ATP recycling system was performed in presence of 2 mM ATP, 10 mM PEP, 20 mM MgCl_2_, 50 mM KCl, 8 µg/ml *Sso*PK (Haferkamp, Tjaden et al. 2019) with 20 µg/ml of heteromeric *Sa*PPK3 or 20 µg/ml of homomeric cPPK3 (Saci_2019) in 50 mM TRIS/HCl, pH 7.4 with 10 % (v/v) D_2_O (1 ml total volume, 735 min). To follow PEP hydrolysis as well as the effect of *Sa*PPK3 on hydrolysis, a sample, without polyP or ATP, in presence of 10 mM PEP, 20 mM MgCl_2_, 50 mM KCl in 50 mM TRIS/HCl, pH 7.4 with 10 % (v/v) D_2_O was analyzed in presence (1 ml total volume, 810 min) and absence of 20 µg/ml of heteromeric *Sa*PPK3 (1 ml total volume, 750 min). To investigate polyP hydrolysis a sample in presence of 10 mM ADP, 500 µM polyP_45_, 10 mM MgCl_2_, 50 mM KCl, in 50 mM TRIS/HCl, pH 7.4 with 10 % (v/v) D_2_O was analyzed (1 ml total volume, 720 min). The reactions were followed over time at 70 °C, where one merged spectrum was recorded every 15 or 30 min. The signals obtained were assigned (see SI Fig. 11, 12) and integrated over a list of the spectra, where the sum of integrals was normalized.

### Stability of different phosphate compounds at high temperature

To determine the stability of (poly)phosphate compounds at higher temperature in presence and absence of divalent (MgSO_4_) and monovalent ions (KH_2_PO_4_), different reaction mixtures were incubated at 70 °C and qualitative analysis was carried out with 31P NMR spectroscopy. The samples contained 500 µM polyP_45_, 500 µM polyP_45_ with 2.5 mM MgSO_4_, 500 µM polyP_45_ with 500 µM KH_2_PO_4_, or 100 mM cP_3_ in 50 mM TRIS/HCl pH 7.2, 200 mM NH_4_Cl and 10 % D_2_O. The samples were incubated overnight at 70°C the quantification of phosphates was determined by evaluation of ^31^P NMR spectra.

### ^31^P NMR analysis

For detailed analysis of the phosphate compounds, ^31^P NMR analysis was used. Therefore, ^1^H- and ^31^P NMR spectra were recorded on an DRX 300, DRX 500 or AVANCE NEO 400 MHz nuclear magnetic resonance spectrometer (Bruker, Billerica, USA) at ambient temperature in 90 % H_2_O and 10 % D_2_O (for NMR lock), where ^1^H-NMR spectra were recorded only for NMR lock. The ^31^P NMR chemical shifts are given in ppm (for AVANCE NEO 400 MHz); splitting patterns are given as singlet (s), doublet (d), triplet (t), plus coupling constants (J) are reported in Hz. The spectra were analyzed with TopSpin Software (Bruker, Billerica, USA). The NMR spectra are shown in the supplemental material (SI). The spectra were integrated using the multiple method for a list of spectra with normalization setting the sum of integrals to 1 for each spectrum. Since the relaxation time was not determined, the concentrations were not recalculated. Starting from the integrals obtained, the relative molar ratio was calculated by dividing the integrals by the number of phosphor atoms in the molecules giving the respective signal (e.g. three P atoms in cP_3_). For each compound standard ^31^P NMR was performed, revealing the signals summarized in the SI part.

### Model construction

A kinetic model for polyP metabolism was constructed based on initial rate kinetics for the *Sa*PKK3, and degradation kinetics for PolyP and PEP determined via ^31^P NMR. The model had to be adapted in several ways to simulate the polyP dynamics in the incubation assays. These adaptations and the full model description are given here.

In numerous assays (e.g. Fig. 3A, B, C) we observed that independent of the starting conditions, in the presence of polyP, ATP and ADP, an equilibrium ATP/ADP ratio of 3-4 was obtained in PKK3 incubations (similar to what was observed in Keppler et al. 2022 (Keppler, Moser et al. 2022)). Our initial rate kinetics showed a much faster Vm in the reverse (polyP degradation) direction than in the forward direction, and with the measured Km values this would result in a much higher equilibrium ATP/ADP ratio than the observed 3-4. In addition, we observed much faster polyP synthesis rates in the NMR analysis (Fig. 3C) than were observed in the initial rate experiments. To accommodate for these observations, and the gradual slow increase in polyP synthesis when started in absence of a polyP-template (Fig. 3C), we multiplied the Vm forward for the *Sa*PKK3 with a factor (1 + 5 polyP). A second adaptation had to be made to the kinetic constant for the polyP degradation to cP_3_, which was 1.5 times faster in the enzyme incubations than was observed in the isolated polyP degradation. Lastly the uncoupled polyP degradation was observed to be *Sa*PPK3 dependent and had to be multiplied with a factor 4x for 20 ug/ml incubations compared to the 5 ug/ml incubations.

### Bioinformatic analysis and computational analysis

Crystal structures were retrieved from the pdb database as indicated. Structural models were retrieved from the AlphaFold Protein Structure database (Varadi, Anyango et al. 2021) or predicted using the the alphafold server (Alphafold 3 (Abramson, Adler et al. 2024)). SwissDock was used to infere the putative polyP binding sites in Saci_2019 (Grosdidier, Zoete et al. 2011) and poly9P was used as ligand available in the pdb database (9PI). Two of the best hits overlapping in the putative binding cleft were further used and manually adjusted to predict the position of a polyP stretch comprising 12 phosphate moieties. Structural analyses, comparisons, and visualizations were done using UCSF ChimeraX package from the Resource for Biocomputing, Visualization, and Informatics at the University of California, San Francisco, with support from National Institutes of Health R01-GM129325 and the Office of Cyber Infrastructure and Computational Biology, National Institute of Allergy and Infectious Diseases (Meng, Goddard et al. 2023). Alignments were performed with clustal omega using the geneious prime software (Geneious Prime (https://www.geneious.com) or EMBL server (Lombard, Camon et al. 2002). The alignments were then used in the MEGA11 software package for phylogenetic tree constructions (Tamura, Stecher et al. 2021) (for further details see legend to Figure 6).

## Supporting information

Supplementary Material_revised

## Abbreviations

polyP: polyphosphate
VTC: complex vacuolar transporter chaperone complex
PPK: polyphosphate kinase
PPN: endopolyphosphatase
PPX: exopolyphosphatase
dTMPK: thymidylate kinase
dTMP: deoxythymidine monophosphate
dTDP: deoxythymidine diphosphate

