## Supplementary Material_revised for "The archaeal family 3 polyphosphate kinase reveals a function of polyphosphate as energy buffer under low energy charge"

### Supplementary Figures

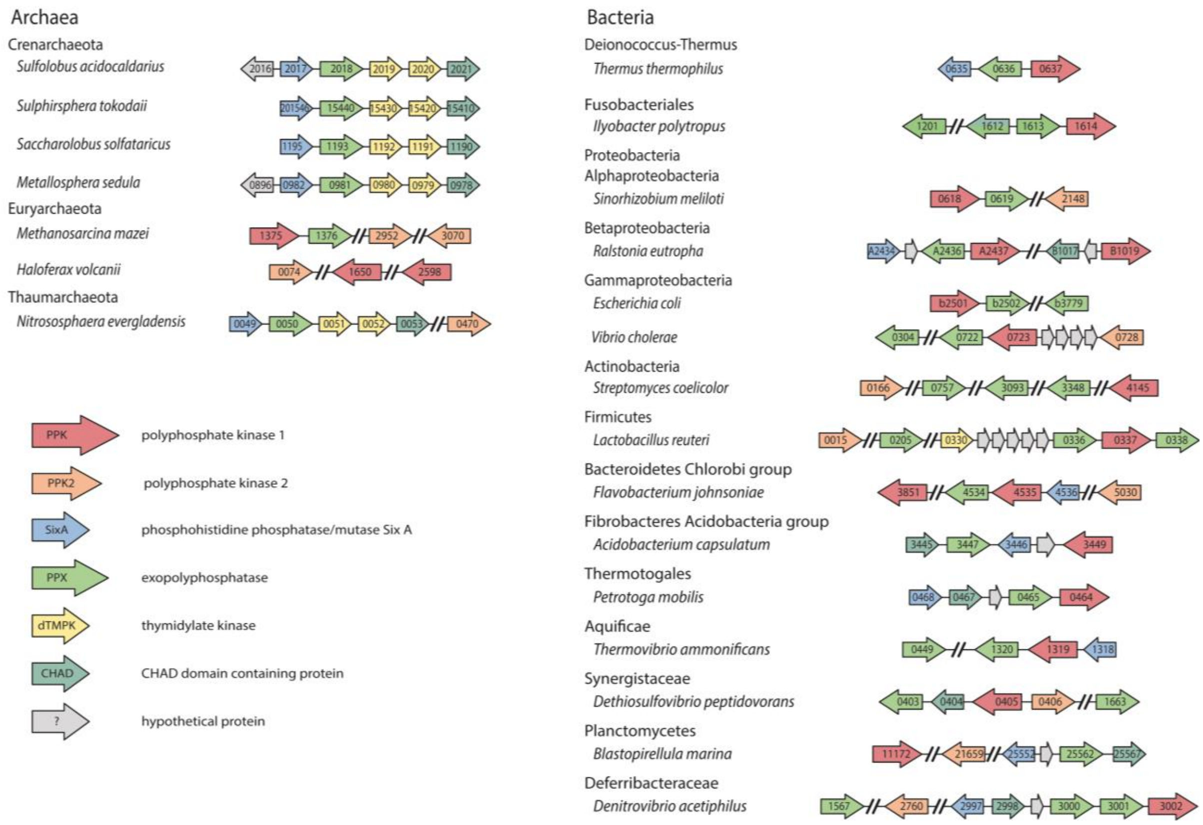


**SI Figure 1: Gene neighborhood analysis of polyP metabolizing enzymes in bacteria and archaea.** The operon and neighborhood relations were investigated using the KEGG and STRING databases. The scheme shows genes encoding the polyphosphate kinase family 1 (red), 2 (orange), the phosphohistidine phosphatase/mutase SixA (blue), the exopolyphosphatase (green), the thymidylate kinase (yellow), the CHAD domain containing protein (turquoise) and hypothetical proteins (grey). An arrow in the respective color represents each gene with the associated gene number.

Among bacteria and some archaeal species, the *ppk* gene is occurring in the immediate neighborhood of the *ppx* gene. Only in crenarchaeal organisms, the gene organization did not present a PPK homolog. However, the *ppx* gene cluster within Crenarchaeota seems to be conserved, containing *sixA, ppx,* two *dtmpk* and *chad* genes, thus suggesting one of these genes has an effect on polyP metabolism.

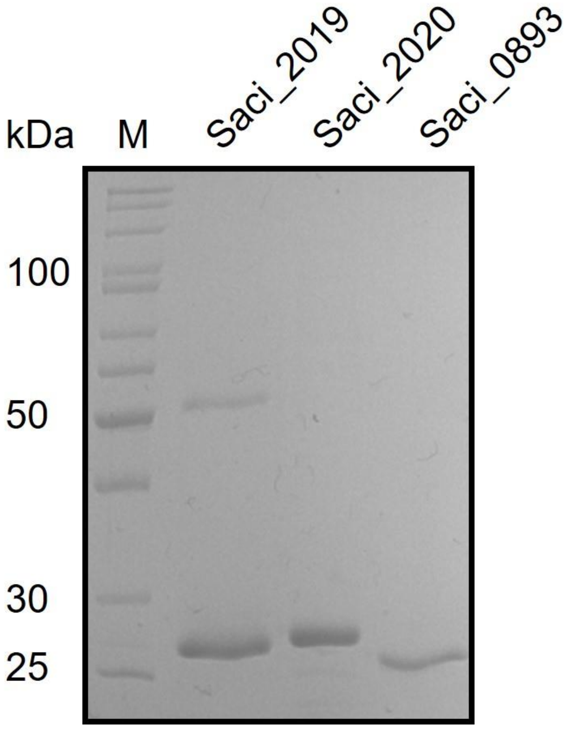


**SI Figure 2: The three annotated dTMPKs (Saci_2019, Saci_2020 and Saci_0893) from *S. acidocaldarius* after recombinant expression in *E. coli* Rosetta (DE3) and purification via affinity chromatography (Ni-TED column) (SDS-PAGE and Coomassie staining).** The two subunits of PPK dTMPK_2,3, (*saci_2019, saci_2020*) and thymidylate kinase dTMPK_1 (s*aci_0893).* For all enzymes 2 µg of protein were applied. M, protein marker (PageRuler Unstained Protein Ladder 10-200 kDa, Thermo Fisher Scientific, Schwerte, Germany). The calculated molecular masses are 22.58 kDa for dTMPK_1, 24.64 kDa for dTMPK_2 and 23.83 kDa for dTMPK_3, respectively.

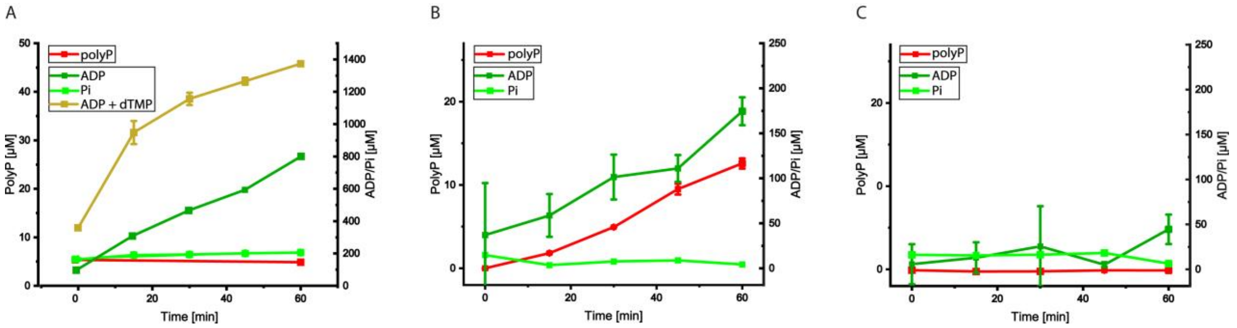


**SI Figure 3: Enzymatic activity of the three thymidylate kinase homologs Saci_0893 (dTMPK_1), Saci_2019 (dTMPK_2) and Saci_2020 (dTMPK3).** The dTMPK assay for Saci_0893 (A), Saci_2019 (B) and Saci_2020 (C) was performed in presence of 2 mM ATP, 50 mM KCl, 4 mM $MgCl_2$ and 10 µg/ml enzyme in 50 mM TRIS/HCl, pH 7.4 following ATP-dependent formation of polyP, ADP, and $P_i$ over time using the polyp assay, the PK-LDH assay, and the malachite green assay, respectively. Since no increase in ADP formation was observed for Saci_2019 and Saci_2020 in presence of dTMP only for Saci_0893 the additional ADP formation with dTMP (1 mM) is shown. Only for Saci_0893 the enzymatic activity was dependent on dTMP as shown by the increased ADP-formation in presence of dTMP. No polyP formation is observed (A). For Saci_2019 ADP as well as polyP formation were detected (B), whereas Saci_2020 showed neiter ADP or polyP formation (C). For all experiments three independent measurements (n=3) were performed and error bars indicate the standard error of the mean (SEM).

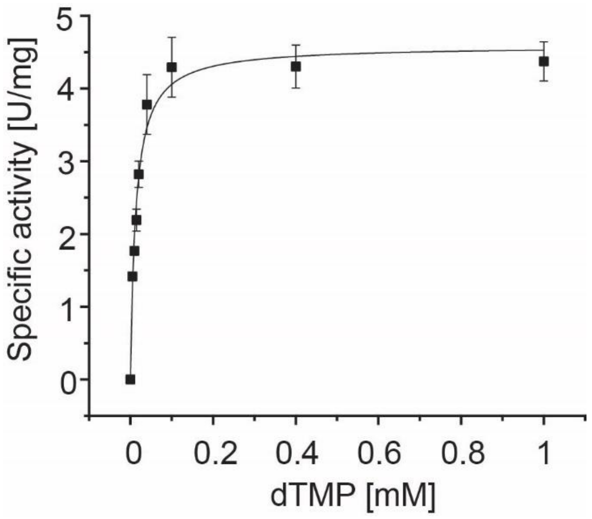


**SI Figure 4 Characterization of recombinant dTMPK (Saci_0893)**. The kinetic properties were determined at 55°C in the presence of 0.05 – 1 mM dTMP, 2 mM ATP, 4 mM $MgCl_2$, 0.2 mM NADH and 1.25 µg/mL Saci_0893 in 50 mM TRIS/HCl, pH 7.4 coupled with PK-LDH (0.5 ml total volume). For all experiments three independent measurements (n=3) were performed and error bars indicate the standard error of the mean (SEM).

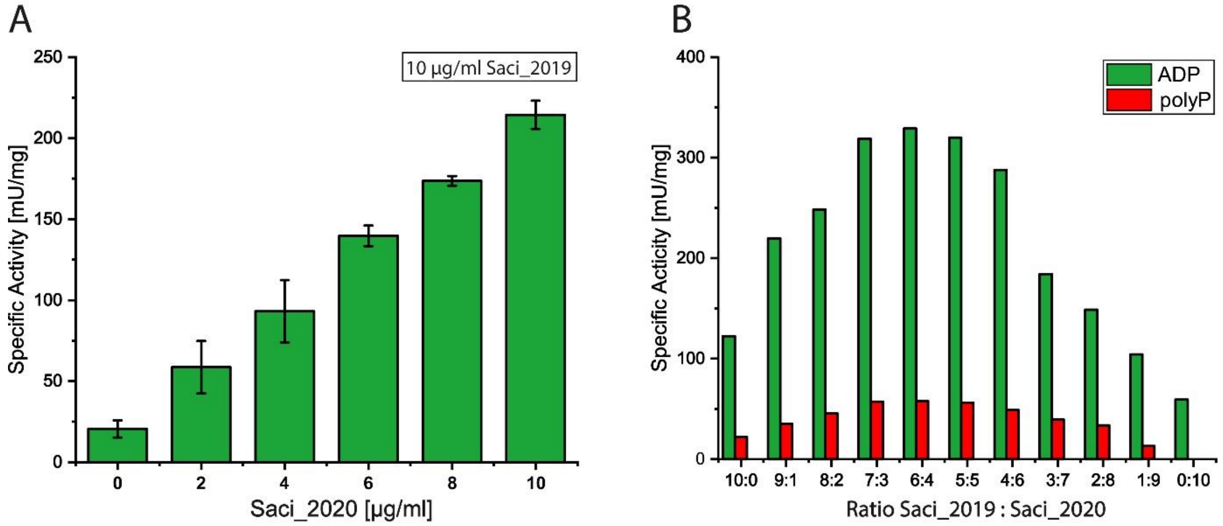


**SI Figure 5: Titration experiments to address the activation of homomeric PPK (Saci_2019) by Saci_2020.** (A) The specific PPK activity of homomeric PPK (Saci_2019, 10 µg/ml protein) upon addition of 0 - 10 µg/ml Saci_2020 (0.5 ml total volume) was determined by following ATP-dependent ADP formation over time at 70°C using the PK-LDH assay. (B) The formation of ADP (green) and polyP (red) was determined (using the PK-LDH and polyp assay, respectively) in presence of different molar ratios of Saci_2019 and Saci_2020 (from 10:0 to 0:10, total enzyme concentration 20 µg/ml).

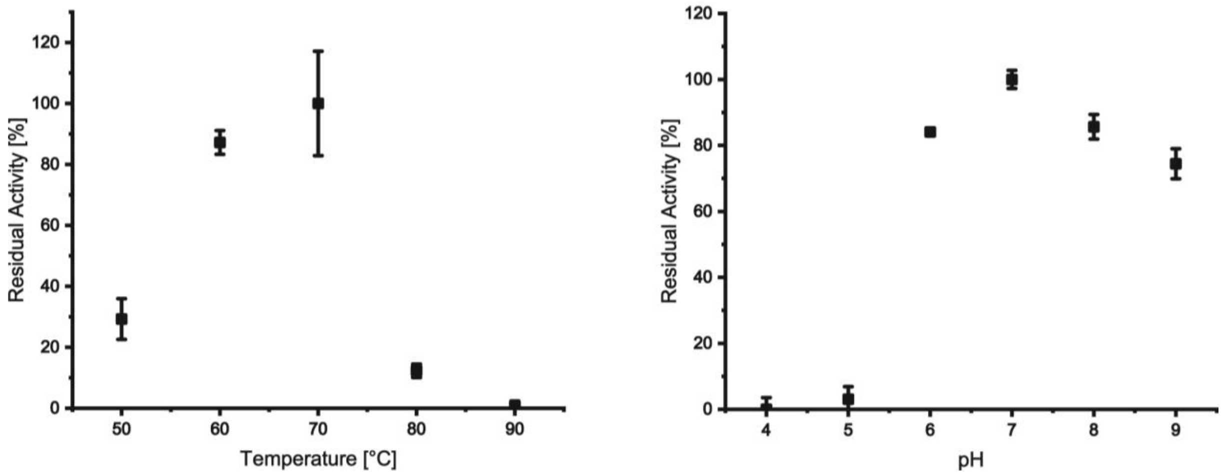


**SI Figure 6: Temperature and pH optimum of the heteromeric *Sa*PPK3 (Saci_2019 and Saci_2020 (1:1)).** The temperature (A) and pH (B) dependence was determined in the direction of polyP formation. The enzymatic activity was determined at 70°C using 20 µg/ml of *Sa*PPK3 by detecting ADP formation (discontinuous PK-LDH assay). The temperature dependence (50 – 90 °C) was analyzed in 50 mM TRIS/HCl, pH 7.4 (adjusted at the respective temperature) and the pH dependence in a mixed buffer system (50 mM TRIS/HCl, Citrate/HCl, BICINE/NaOH and MES/NaOH, respectively) adjusted to the respective pH (pH range 4-9) at 70°C. For all experiments three independent measurements (n=3) were performed and error bars indicate the standard error of the mean (SEM).

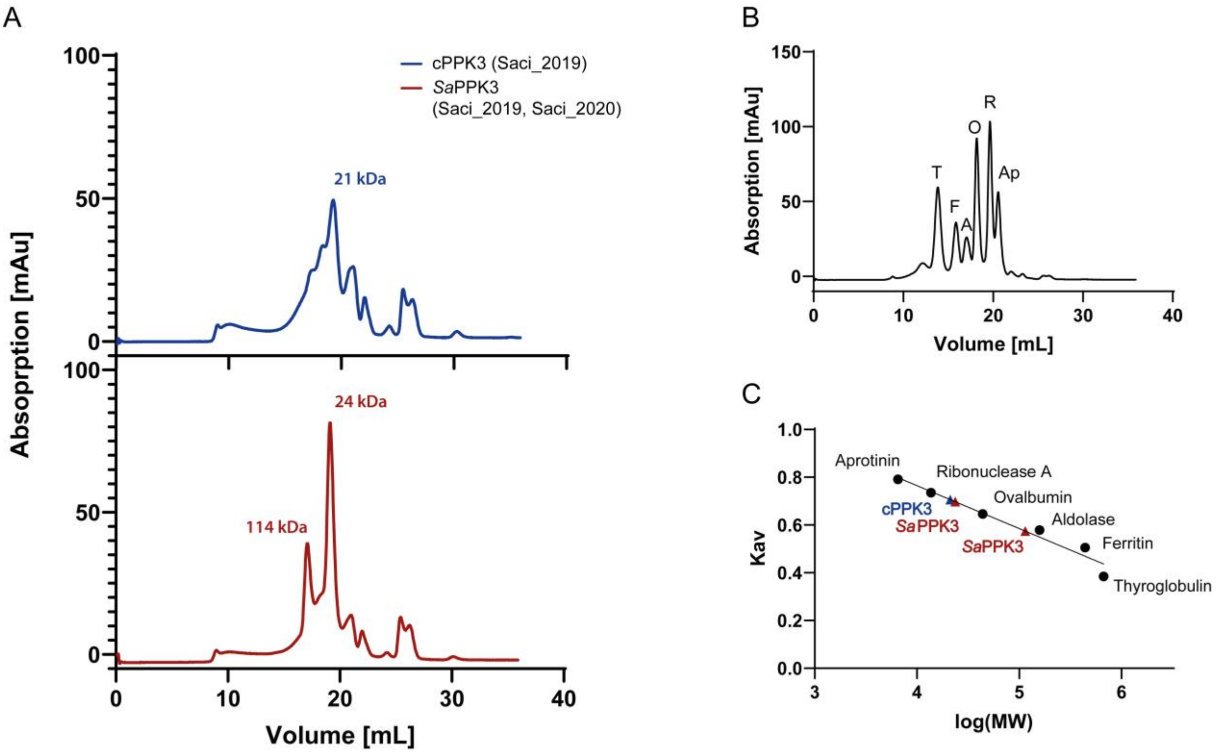


**SI Figure 7: Native molecular mass of cPPK3 (Saci_2019) and heteromeric *Sa*PPK3.** (A) Size exclusion chromatography (Superose® 6 Increase 10/300 GL, Cytiva Lifescience™, Freiburg, Germany) of the homomeric cPPK3 (Saci_2019) and heteromeric *Sa*PPK3 (Saci_2019 and Saci_2020 (1:1)) from *S. acidocaldarius*. (B, C) Protein standards (aprotinin (6.5 kDa), ribonuclease (13.7 kDa), ovalbumin (44 kDa), aldolase (158 kDa), ferritin (440 kDa) and thyroglobulin (669 kDa)) from the LMW and HMW gel filtration calibration kits (Cytiva Lifescience™) were applied to generate a calibration curve for determination of the molecular masses of the recombinant enzymes. The elution profile of the homomeric cPPK3 (blue) shows a monomeric structure with higher aggregation states. In presence of the second subunit (heteromeric *Sa*PPK3, red) a shift in the molecular mass towards tetramer formation is observed, however, monomers are still present.

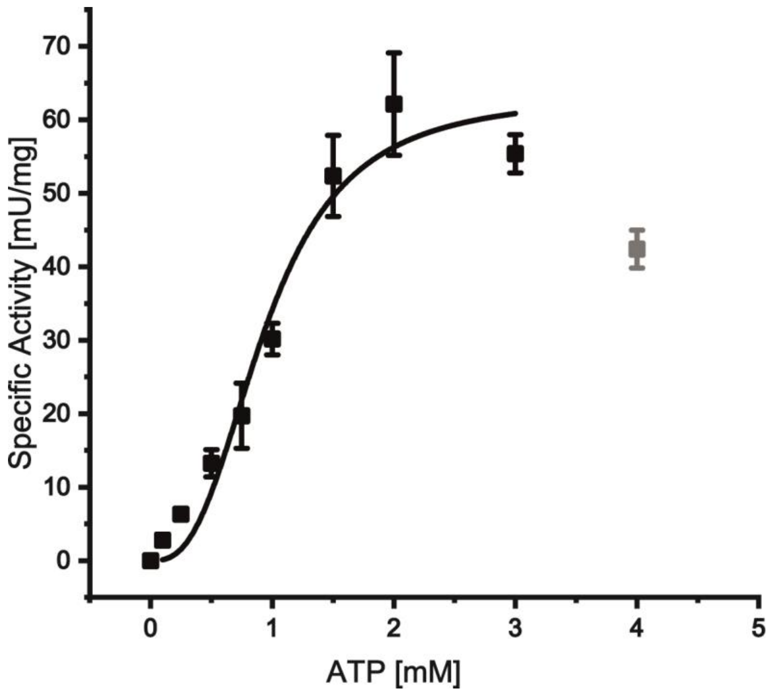


**SI Figure 8: Kinetic characterization of the recombinant heteromeric *Sa*PPK3 (Saci_2019 and Saci_2020 (1:1)) determined following polyP formation.** The kinetic properties were determined at 70°C in the presence of 0 – 4 mM ATP, 4 mM $MgCl_2$, 50 mM KCl and 10 µg/ml of *Sa*PKK3 in 50 mM TRIS/HCl, pH 7.4 using the discontinuous polyP assay. For all experiments three independent measurements (n=3) were performed and error bars indicate the standard error of the mean (SEM).

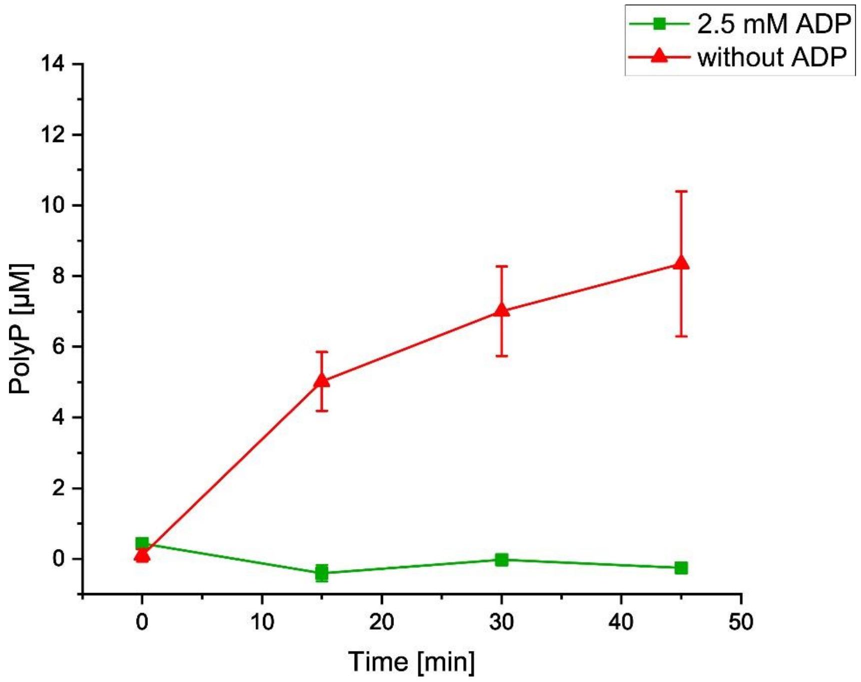


**SI Figure 9: Inhibitory effect of ADP on polyP formation by the heteromeric *Sa*PPK3.** The polyP formation was followed during PPK assay (as described) in absence as well as presence of 2.5 mM ADP to the reaction mixture containing 1.5 mM ATP as substrate. The polyP formation at 70°C was followed using polyP assay. For all experiments three independent measurements (n=3) were performed and error bars indicate the standard error of the mean (SEM).

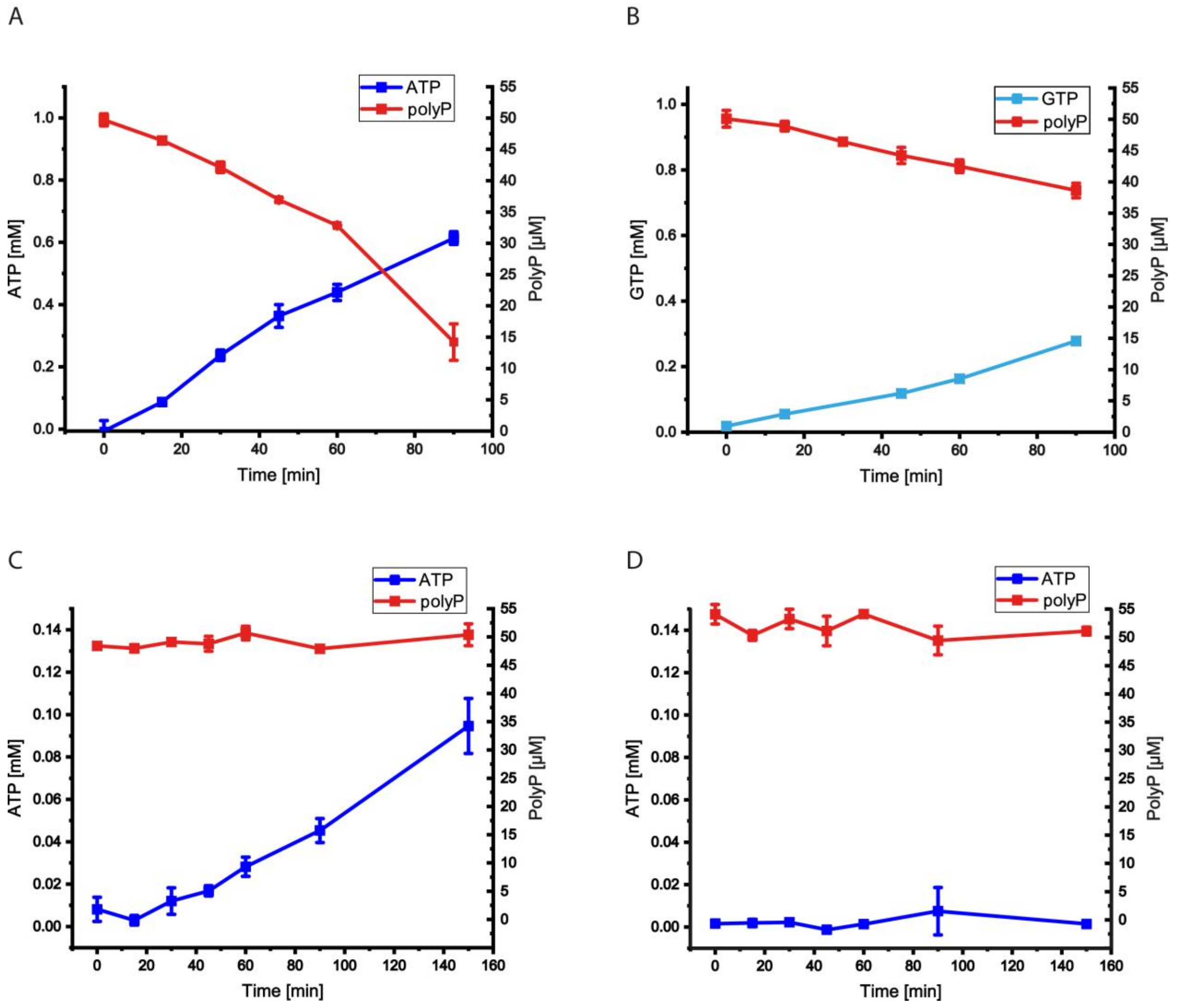


**SI Figure 10: PolyP-dependent nucleotide kinase activity of the recombinant heteromeric *Sa*PPK3 and the homomeric PPKs (Saci_2019 and Saci_2020) from *S. acidocaldarius* with respect to ATP and GTP formation.** The polyP-dependent formation of ATP from ADP (A) and GTP from GDP (B) with 5 µg/ml of heteromeric *Sa*PPK3 as well as the ATP formation with 5 µg/ml of the homomeric enzymes Saci_2019 (C) and Saci_2020 (D) at 70°C are shown. The enzymatic activity (i.e. ATP/GTP formation and polyP utilization) was determined in presence of 50 µM $polyP_{45}$ as phosphoryl donor and 2 mM ADP (A, C, D) or GDP (B) as phosphoryl acceptor at 70 °C in 10 mM $MgCl_2$, 50 mM KCl, 50 µM $polyP_{45}$ in 50 mM TRIS/HCl, pH 7.4. The ATP (blue squares) or GTP formation (light blue squares) was determined by using the discontinuous HK-G6PDH assay and polyP degradation was quantified via the polyP assay. For all experiments three independent measurements (n=3) were performed and error bars indicate the standard error of the mean (SEM).

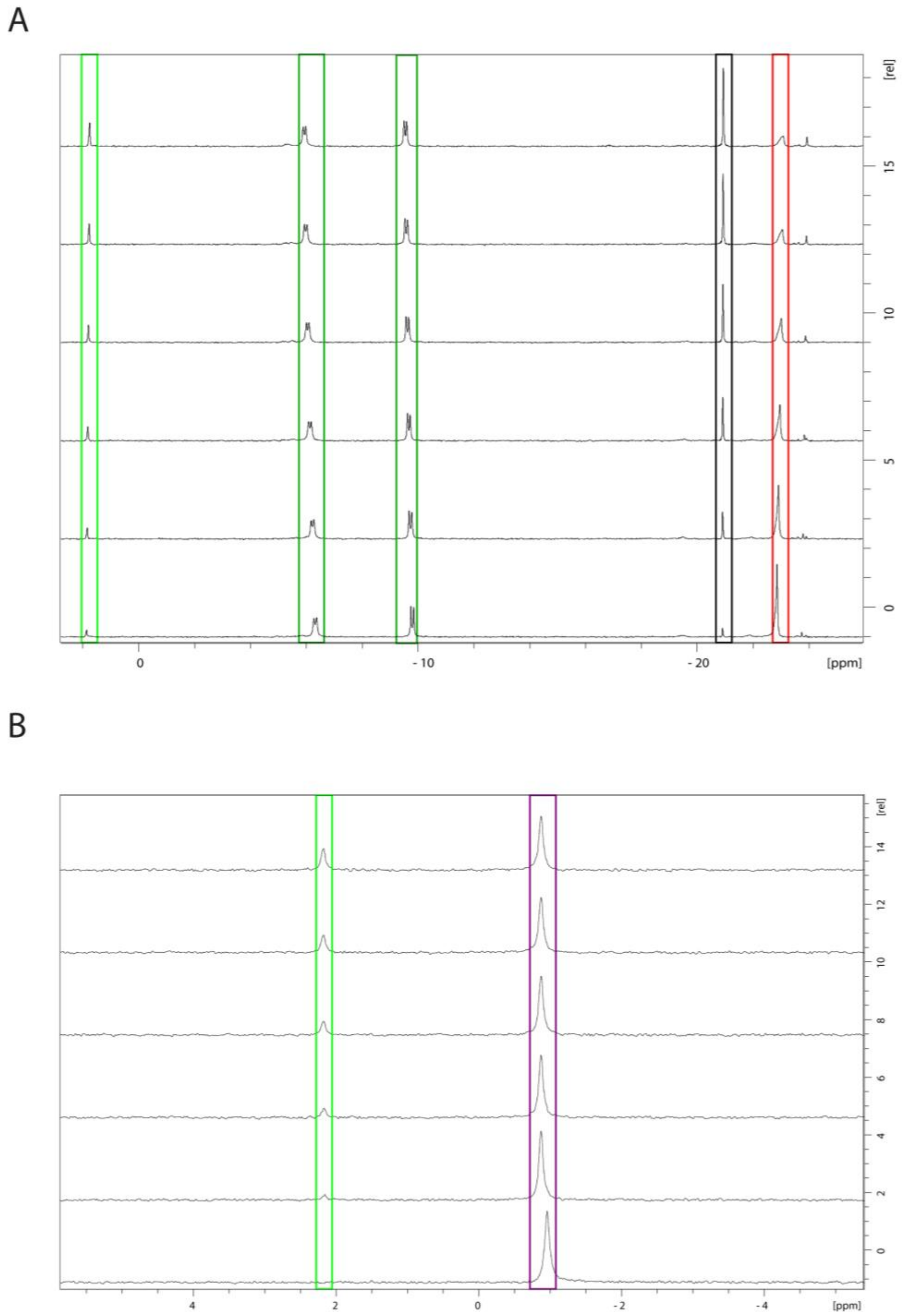


**SI Figure 11: Stacked $^{31}P$ NMR spectra of non-enzymatic degradation of polyP, and phosphoenolpyruvate (PEP) hydrolysis (SI Fig. 13)**. All assays were performed at 70°C in 50 mM TRIS/HCl pH, 7.4 with 50 mM KCl in 10% (v/v) $D_2O$ (1 mL total volume). (A) The non-enzymatic hydrolysis of polyP to $cP_3$ and $P_i$ was analyzed in presence of 20 mM ADP, 500 µM $polyP_{45}$, 10 mM $MgCl_2$. (B) The non-enzymatic $P_i$ liberation from PEP was performed with 20 mM $MgCl_2$, 10 mM PEP in presence of 20 µg/ml heteromeric *Sa*PPK3. The obtained signals in the $^{31}P$ NMR spectra were identified as ADP (dark green), polyP (red), $P_i$ (light green), PEP (purple) and $cP_3$ (black). The reaction was followed from 0 min (bottom spectrum) to 720 (A) or 810 min (B) (top spectrum), respectively.

A

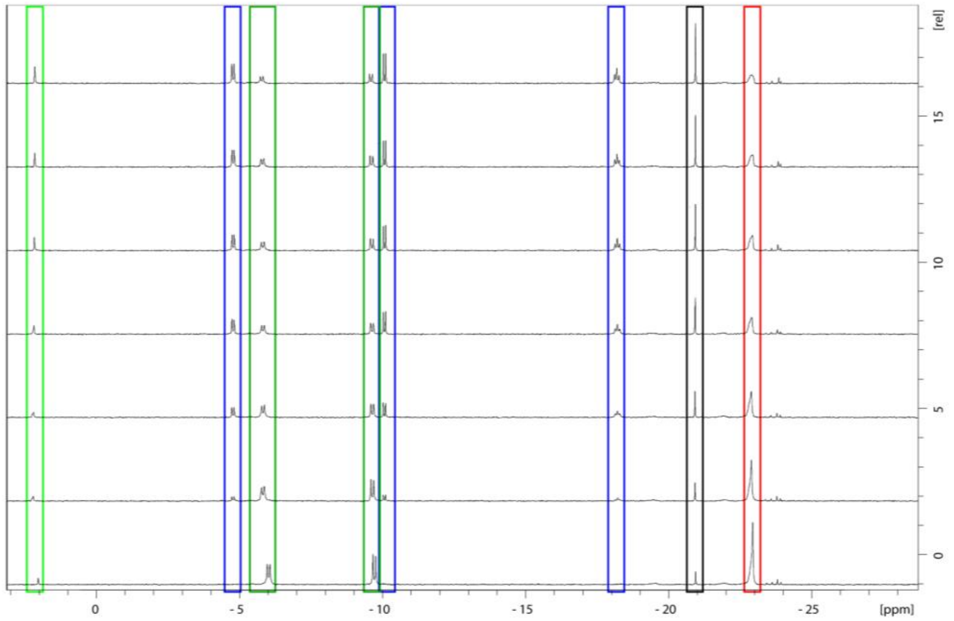


B

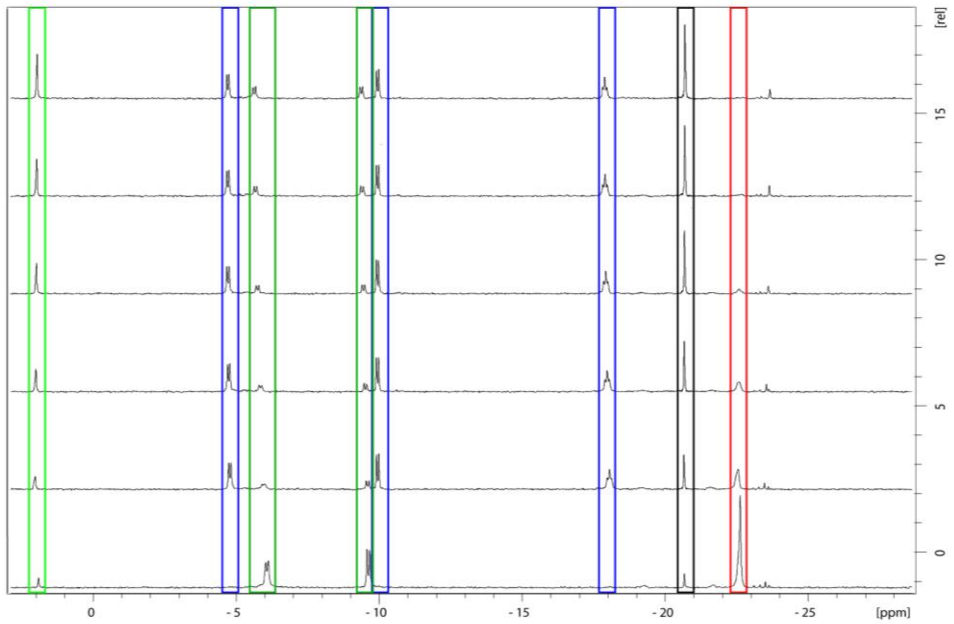

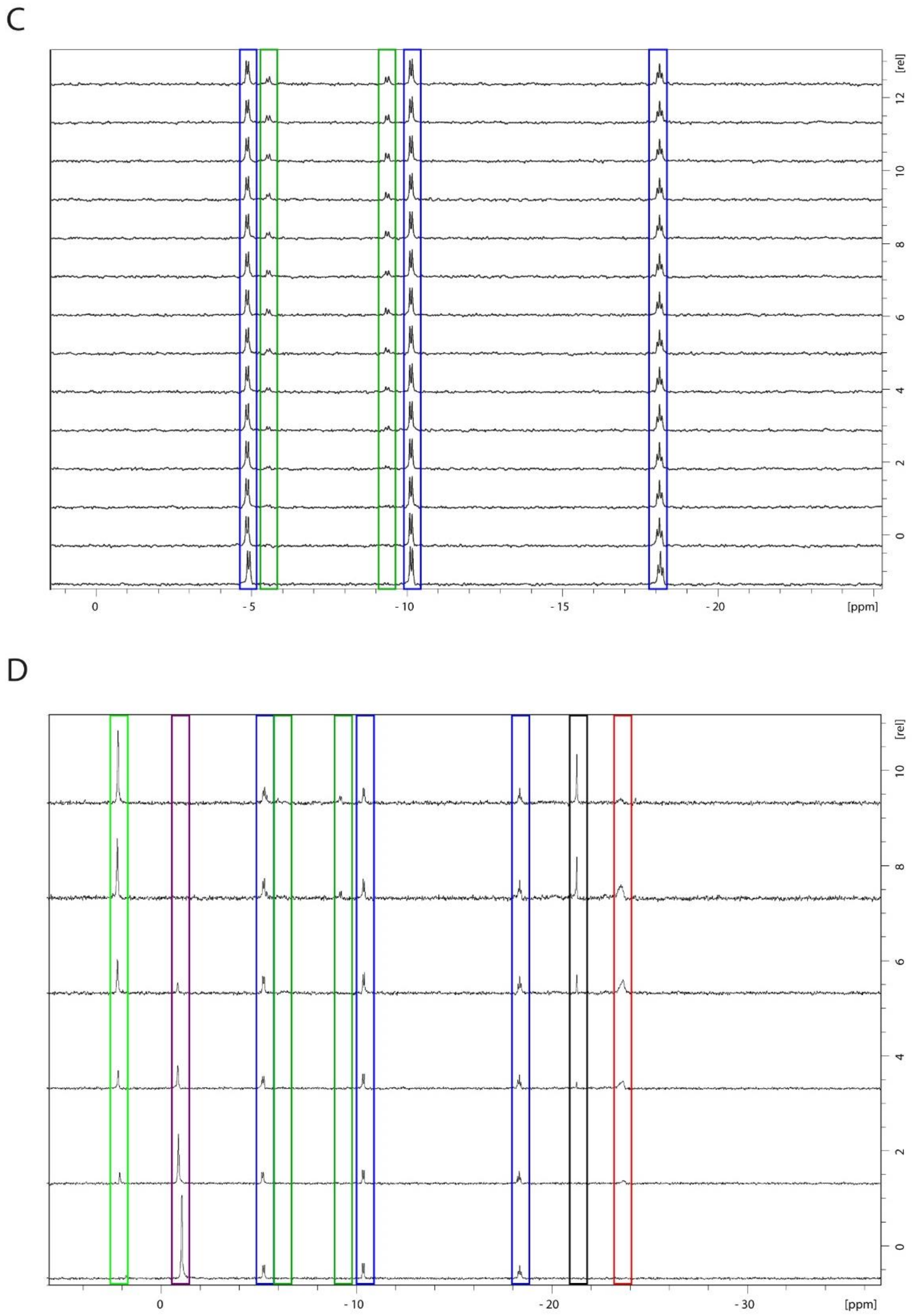


**SI Figure 12: Stacked $^{31}$P NMR spectra of the reversible *Sa*PPK3 activity in the direction of NTP (A, B) and polyP formation without (C) and with ATP recycling system (D)** All assays were performed in 50 mM TRIS/HCl pH, 7.4 with 50 mM KCl in 10% (v/v) $D_2O$ (1 mL total volume). The polyP-dependent ATP formation was performed with 20 mM ADP, 500 µM $polyP_{45}$, 10 mM $MgCl_2$ with 5 µg/ml (A) and 20 µg/ml (B) heteromeric *Sa*PPK3. The ATP-dependent polyP formation was performed with 2 mM ATP and 4 mM $MgCl_2$ with 20 µg/ml heteromeric PPK without (C) and with ATP recycling (20 mM $MgCl_2$, 10 mM PEP, 8 µg/mg *Sso*PK) (D). The signals obtained in the $^{31}$P NMR spectra were identified as polyP (red), $cP_3$ (black), ATP (blue), ADP (dark green), PEP (purple), $P_i$ (light green). The reaction was followed at 70°C from 0 min (bottom spectrum) to 300 min, 750 min, 360 min, 735 min (top spectrum), respectively.

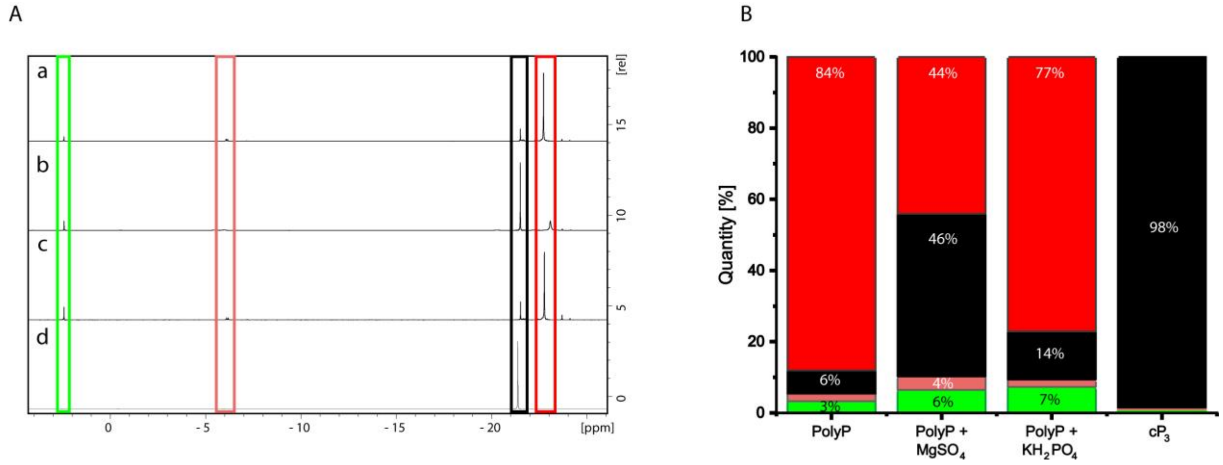


**SI Figure 13: Stability of different phosphate compounds in presence and absence of divalent ($MgSO_4$) and monovalent ($KH_2PO_4$) ions analyzed via $^{31}$P NMR.** (A) NMR spectra of samples incubated at 70°C overnight in 50 mM TRIS/HCl pH 7.2, in presence of 200 mM $NH_4Cl$ with 500 µM polyP$_{45}$ (a), 500 µM polyP$_{45}$ with 2.5 mM $MgSO_4$ (b), 500 µM polyP$_{45}$ with 500 µM $KH_2PO_4$ (c) and 100 mM cP$_3$ (d). (B) The quantity [%] of the different phosphate compounds with P$_i$ (green), polyP$_3$ (light red), cP$_3$ (black) and polyP (red) is shown.

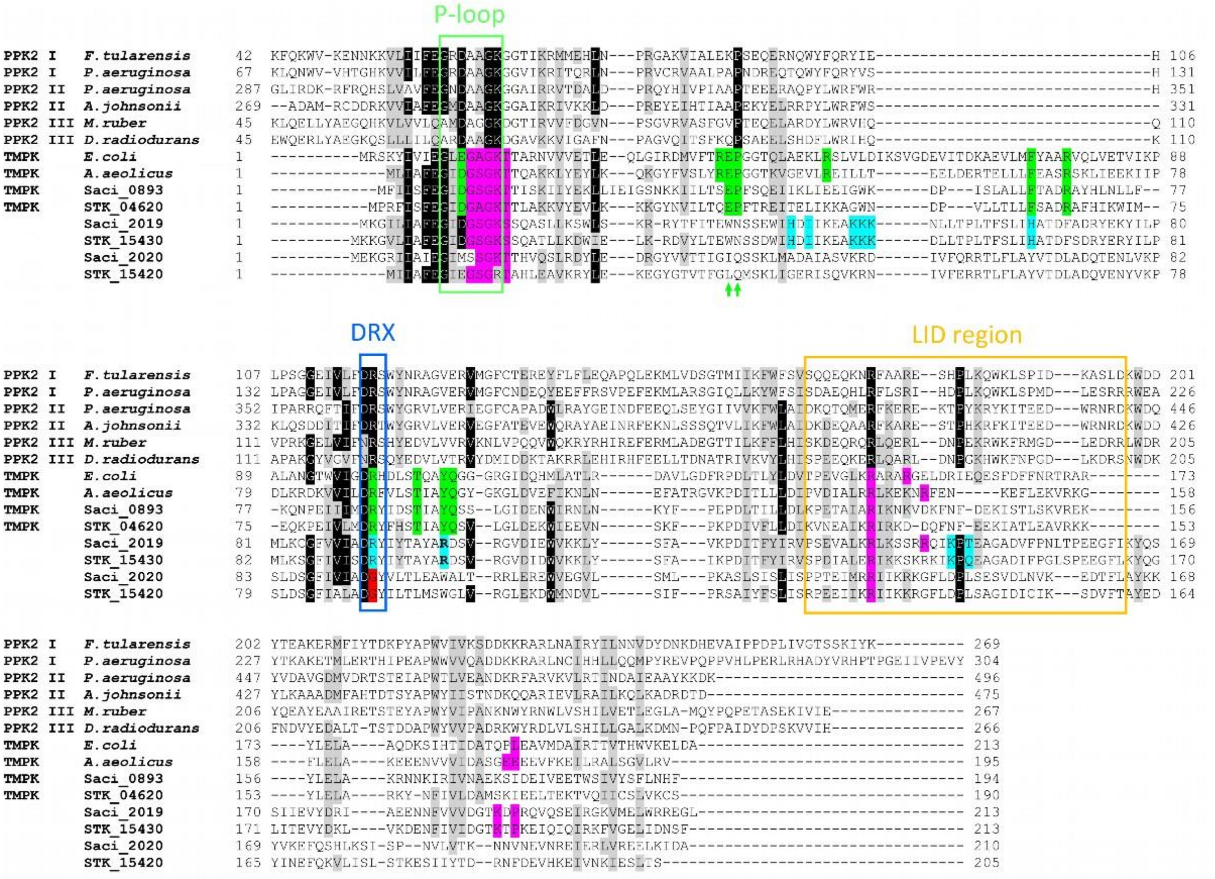


**SI Figure 14**. **Sequence alignment of** ***the S. acidocaldarius*** **Saci_2019 cPPK3 and Saci_2020 rPPK3 with PPK2s and dTMPKs**. The alignment was generated by clustal omega (Sievers, Wilm et al. 2011, Madeira, Madhusoodanan et al. 2024). The general motifs present in the P-loop kinase superfamily are boxed in green (P-loop, Walker A), blue (DRX (Walker B) motif) and orange (Lid region) (Leipe, Koonin et al. 2003, Biswas, Shukla et al. 2017). The residues involved in ATP/ADP (phosphate donor) binding (as shown in e.g. the crystal structures from *A. aeolicus* and *E. coli* TMPKs (Lavie, Ostermann et al. 1998, Biswas, Shukla et al. 2017) as well as those predicted from structural models of the cPPK3s are highlighted in magenta. Those of these residues also conserved in PPK2s are mostly involved in interactions with (poly)phosphate moieties (Neville, Roberge et al. 2022). Those residues involved in TMP binding (as shown in the crystal structures from *A. aeolicus* and *E. coli* TMPKs (Lavie, Ostermann et al. 1998, Biswas, Shukla et al. 2017)) including the EP motif (green arrows) (Biswas, Shukla et al. 2017) and their conservation in the archaeal TMPKs are highlighted in green clearly demonstrating that they are not conserved in PPK3s. The residues proposed to form the polyphosphate binding cleft in cPPK3s (Saci_2019 and STK_15430) are shown in cyan and are clearly not conserved in rPPK3s (Saci_2020 and STK_15420). Also, the catalytically essential arginine residue from the DRX motif functioning as clamp in binding and correct positioning of the terminal phosphate moieties from donor and acceptor (Ostermann, Schlichting et al. 2000, Biswas, Shukla et al. 2017) is clearly not conserved in rPPK3s (highlighted in red) explaining its missing activity. For clarity the N-terminal extensions present in PPK2s are not shown.

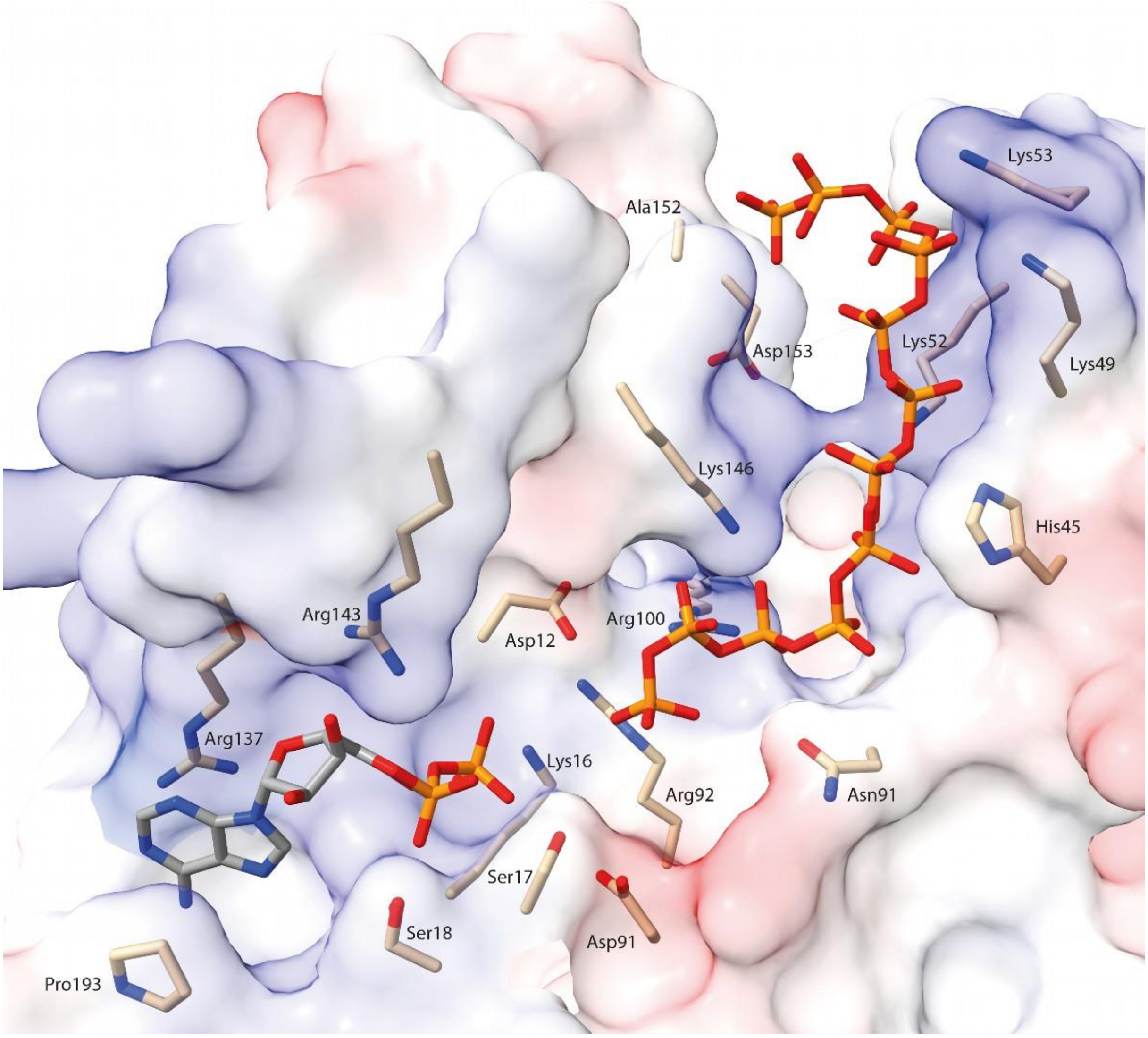


**SI Figure 15. Detailed close-up view of the ADP and polyP binding site proposed for the Saci_2019 cPPK3.** ADP is shown in the same position as in the crystal structure of *S. tokodaii* STK_15430. The proposed polyphosphate binding site in the Saci_2019 cPPK3 (A) (polyphosphate shown as stick model) was identified by swissdock (Grosdidier, Zoete et al. 2011, Bugnon, Röhrig et al. 2024) based docking experiments using the polyP$_9$ available in the pdb database as ligand. Two of the best hits are shown as stick models (with overlapping phosphate moieties after manual adjustments). Saci_2019 is shown as transparent surface representation (blue, positive charge; red negative charge). ADP, polyphosphate, and the residues involved in ADP and polyP binding are shown as stick models and amino acids are labelled. Together with the alignment in SI Fig. 14 it becomes evident that both binding sites are highly conserved in cPPK3 whereas the polyP binding site appears to be absent from rPPK3s.

**Supplementary Tables**

**SI Table 1: List of primers for cloning, plasmids used and constructed in this work, and strains employed**

| **Primers** | **Sequence (5’-3’)** | |
|---|---|---|
| ***saci_0893* fwd *Nde*I** | GGCGCCATATGTTTATAATATCATTTGAAGGAATAGATGG | |
| ***saci_0893* rev *Xho*I** | GCGGCCTCGAGTTAGAAATGGTTTAGAAAAGAATAAAC | |
| ***saci_2018 ppx* fwd *Nde*I** | CCCCATATGCGATATGCGGTAATAG | |
| ***saci_2018 ppx* rev *Xho*I** | GCGCCTCGAGGTCATAGCACACCAGCCACTGAC | |
| ***saci_2019 dtmpk* rev *Nco*I** | GCGCCATGGTGCCGTCAACAACAACAAAG | |
| ***saci_2019 dtmpk* fwd *BamH*I** | GCCGGATCCATATAAGGGCAATAGCGAAG | |
| ***saci_2020 dtmpk* fwd *Nde*I** | CCCCCATATGGATGGAGAAGGGAAGGATTATAGC | |
| ***saci_2020 dtmpk* fwd *BamH*I** | CCCCGGATCCCCTCAGGCGTCAATTTTCAGCTC | |
| **Plasmids** | | |
| **pET15b** | *E.coli* expression plasmid carrying an N-terminal His Tag | Novagen, USA |
| **pET28b** | *E.coli* expression plasmid carrying an C- or N-terminal His-Tag | Novagen, USA |
| **pET15b-*Saci_0893*** | *E. coli* expression plasmid of *dtmpk* (*saci_0893*) cloned into pET15b | This work |
| **pET15b-*Saci_2018*** | *E. coli* expression plasmid of *ppx* (*saci_2018*) cloned into pET15b | This work |
| **pET15b-*Saci_2019*** | *E. coli* expression plasmid of *dtmpk* (*saci_2019*) cloned into pET15b | This work |
| **pET28b-*Saci_2020*** | *E. coli* expression plasmid of *dtmpk* (*saci_2020*) cloned into pET28b | This work |
| **Strains** | **Source/Reference** | |
| ***E. coli* DH5□** | Hanahan, USA | |
| ***E. coli* Rosetta (DE3)** | Stratagene, USA | |

**NMR spectroscopy**: $^{1}$H- and $^{31}$P NMR spectra were recorded on an DRX 300, DRX 500 or AVANCE NEO 400 MHz nuclear magnetic resonance spectrometer (Bruker, Billerica, USA) at ambient temperature in 90 % $H_2O$ and 10 % $D_2O$. The $^{31}$P NMR chemical shifts are given in ppm (for AVANCE NEO 400 MHz); splitting patterns are given as singlet (s), doublet (d), triplet (t), plus coupling constants (J) are reported in Hz.

polyP$_n$

$^{31}$P NMR(400MHz, 90 % $H_2O$ and 10 % $D_2O$) δ[ppm] = -0.71 (s, 2P, 1-P; n-P), -8.43 (t, $^{2}J_{2,1}$=14.34 Hz, 2P, 2-P; (n-2)-P), -21.67 (s, (n-4)P, (3 to (n-2))-P).

P$_3$

$^{31}$P NMR(400MHz, 90 % $H_2O$ and 10 % $D_2O$) δ[ppm] = -5.41 (d, $^{2}J_{1,3}$=19.4 Hz , 2P, 1-P, 3-P), -20.15 (t, $^{2}J_{2,1}$=19.4 Hz, 1P, 2-P).

cP$_3$

$^{31}$P NMR(400MHz, 90 % $H_2O$ and 10 % $D_2O$) δ[ppm] = -20.15 (s, 1P, 1-P, 2-P, 3-P).

PP$_i$

$^{31}$P NMR(400MHz, 90 % $H_2O$ and 10 % $D_2O$) δ[ppm] = -6.00 (s, 2P, 1-P, 2-P).

P$_i$

$^{31}$P NMR(400MHz, 90 % $H_2O$ and 10 % $D_2O$) δ[ppm] = 2.42 (s, 1P, 1-P).

ATP

$^{31}$P NMR(400MHz, 90 % $H_2O$ and 10 % $D_2O$) δ[ppm] = -5.47 (d, $^{2}J_{□□□□}$=15.2 Hz 1P, □-P), -10.63 (d, $^{2}J_{□□□}$=15.2 Hz, 1P, □-P), -19.05 (t, 1P, □-P).

ADP

$^{31}$P NMR(400MHz, 90 % $H_2O$ and 10 % $D_2O$) δ[ppm] = -6.08. (d, $^{2}J_{□,□}$=18.5 Hz, 1P, □-P), -10.14. (d, $^{2}J_{□□□}$=18.5 Hz, 1P, □-P).

ADP

$^{31}$P NMR(400MHz, 90 % $H_2O$ and 10 % $D_2O$) δ[ppm] = -6.08. (d, $^{2}J_{□,□}$=18.5 Hz, 1P, □-P), -10.14. (d, $^{2}J_{□□□}$=18.5 Hz, 1P, □-P).

AMP

$^{31}$P NMR(400MHz, 90 % $H_2O$ and 10 % $D_2O$) δ[ppm] = 4.20 (s, 1P, □-P)

dTMP

$^{31}$P NMR(400MHz, 90 % $H_2O$ and 10 % $D_2O$) δ[ppm] = 3.52 (s, 1P, □-P)

dTDP

$^{31}$P NMR(400MHz, 90 % $H_2O$ and 10 % $D_2O$) δ[ppm] = -10.09 (d, $^{2}J_{□,□}$=21.2, 1P, □-P), -11.45 (d, $^{2}J_{□,□}$=21.2 Hz, 1P, □-P)

PEP

$^{31}$P NMR(400MHz, 90 % $H_2O$ and 10 % $D_2O$) δ[ppm] = -1.08 (s, 1P, □-P).
